# AHK5 mediates ETR1-initiated multistep phosphorelay in *Arabidopsis*

**DOI:** 10.1101/2021.09.16.460643

**Authors:** Agnieszka Szmitkowska, Abigail Rubiato Cuyacot, Blanka Pekárová, Markéta Žd’árská, Josef Houser, Jan Komárek, Zuzana Jaseňáková, Aswathy Jayasree, Michael Heunemann, Elena Ubogoeva, Ioannis Spyroglou, Martin Trtilek, Victoria Mironova, Klaus Harter, Elena Zemlyanskaya, Lukáš Žídek, Michaela Wimmerová, Jan Hejátko

## Abstract

Plants, like other sessile organisms, need to sense many different signals, and in response to them, modify their developmental programs to be able to survive in a highly changing environment. The multistep phosphorelay (MSP) in plants is a good candidate for a response mechanism that integrates multiple signal types both environmental and intrinsic in origin. Recently, ethylene was shown to control MSP activity via the histidine kinase (HK) activity of ETHYLENE RESPONSE 1 (ETR1)^1,2^, but the underlying molecular mechanism still remains unclear.

Here we show that although ETR1 is an active HK, its receiver domain (ETR1_RD_) is structurally and functionally unable to accept the phosphate from the phosphorylated His in the ETR1 HK domain (ETR1_HK_) to initiate the phosphorelay to ARABIDOPSIS HISTIDINE-CONTAINING PHOSPHOTRANSMITTERs (AHPs), the next link downstream members in MSP signaling. Instead, ETR1 interacts with another HK ARABIDOPSIS HISTIDINE KINASE 5 (AHK5) and transfers the phosphate from ETR1_HK_ through the receiver domain of AHK5 (AHK5_RD_), and subsequently to AHP1, AHP2 and AHP3, independently of the HK activity of AHK5. We show that AHK5 is necessary for ethylene-initiated, but not cytokinin-initiated, MSP signaling *in planta* and that it thus mediates hormonal control of root growth.

## Main

In the MSP cascade, signal-driven autophosphorylation of the sensor HK on its conserved His triggers a downstream His-to-Asp-to-His-to-Asp phosphorelay, phosphorylating AHPs and type B response regulators (B-RRs) that consequently control signal-regulated transcription^3^. The first step in the MSP phosphorelay is an intramolecular transfer of a phosphate moiety from the phospho-His in the HK domain to the Asp of the RD within the sensor HK molecule. Previous studies have shown that ETR1_RD_ is unlikely to be phosphorylated^4^, possibly being involved in phosphorylation-independent signalling^5^. In line with this, we did not detect any evidence of phosphate transfer (phosphorylation of ETR1_RD_ or depletion of the radiolabeled phosphate from phosphorylated ETR1_HK_, see below) after two hours of incubation of autophosphorylated ETR1_HK_ in the presence of ETR1_RD_ (Supplementary Fig. 1A). Something similar, i.e. no detectable phosphotransfer, was observed when an ETR1 fragment comprising both HK and RD domain (ETR1_HK-RD_) was autophosphorylated and used as a possible phosphate donor to AHP proteins (Supplementary Fig. 1B). To examine whether ETR1_RD_ is able to bind divalent cations that are essential for RD activation^5,6^, we conducted titration of ^15^N-labeled ETR1_RD_ with MgCl_2_ or MnCl_2_ and monitored potential Mg^2+^/Mn^2+^ coordination by ETR1_RD_ residues by measuring ^1^H-^15^N heteronuclear single quantum coherence (HSQC) NMR spectra. Comparison of the HSQC spectra of ETR1_RD_ in its free form and in the presence of Mg^2+^ and Mn^2+^ ions (Supplementary Fig. 1C, D) revealed only negligible changes in peak positions (MgCl_2_) and peak intensities (MnCl_2_), indicating that ETR1_RD_ binds neither Mg^2+^ nor Mn^2+^. This was further supported by microscale thermophoresis (MST), where no sign was observed of Mg^2+^ or Mn^2+^ binding to the fluorescently labelled ETR1_RD_ over the range of concentrations tested (up to 50 mM Mg^2+^/Mn^2+^, data not shown). Altogether, these results suggest that ETR1_RD_ is unable to mediate phosphotransfer in ETR1-mediated MSP signaling.

The recently demonstrated ability of ETR1 to control MSP via its HK activity^1,2^ implies that ETR1_HK_ may phosphorylate RD of other receptors active in MSP. To test this possibility, we performed a series of phosphotransfer assays with various RDs from *Arabidopsis* sensor HKs. We selected RD of the well-studied cytokinin receptor AHK4 (AHK4_RD_) and RDs from the two subclades of non-cytokinin HK sensors CKI1 and AHK5 (CKI1_RD_ and AHK5_RD_, respectively), as AHK5_RD_ is the closest paralogue of ETR1_RD_ (Supplemental Fig. 2 and 3). After 30 minutes of incubation, we observed phosphotransfer from ETR1_HK_ (manifested as loss of ETR1_HK_ phosphorylation, see Supplemental Fig. 4A) to all three RDs at all molar ratios tested. As anticipated, presence of higher RD concentrations led to more apparent loss of ETR1_HK_ phosphorylation, suggesting faster phosphotransfer. The most efficient phosphotransfer by far was observed for AHK5_RD_, followed by CKI1_RD_ and AHK4_RD_ (Supplemental Fig. 4A). The phosphorylated AHK5_RD_ signal was not detectable after 30 min of incubation, possibly due to fast RD autodephosphorylation activity driving rapid phosphoaspartate hydrolysis^7^. However, phosphorylated CKI1_RD_ and AHK4_RD_ seem to be more stable as the bands corresponding to the phosphorylated forms of both RDs were visible at all ratios tested. Compared to long (30 min) incubation, phosphorylated AHK5_RD_ was clearly detectable as early as 30 s after addition of AHK5_RD_, as also the decrease in intensity of radiolabeled ETR1_HK_ at the shorter intervals (Fig. 1A). In contrast, CKI1_RD_ and AHK4_RD_ were both phosphorylated only very weakly even after 40 min of incubation at a molar ratio of 1:2 (Fig. 1B, Supplemental Fig. 4B). Quantification of phosphotransfer kinetics from phosphorylated ETR1_HK_ to AHK5_RD_ and CKI1_RD_ (Figs. 1C, D, Supplemental Fig. 5; see also Methods) confirmed that ETR1 exhibits strong kinetic preference for AHK5_RD_ *in vitro*. Additional experiments with ETR1_HK-RD_ as a phosphate donor (Supplemental Figs. 4C, D) demonstrated that the presence of ETR1_RD_ does not prevent the ability of ETR1_HK_ to phosphorylate any of the RDs tested. However, AHK5_RD_ was able to accept the phosphate faster from ETR1_HK_ than from ETR1_HK-RD_ (Fig. 1D). Hence, a possible regulatory role for ETR1_RD_ in ETR1_HK_-mediated phosphorylation cannot be excluded. We can thus conclude that AHK5_RD_ is likely the canonical substrate for the histidine kinase-dependent signaling activity of ETR1.

**Figure 1.**
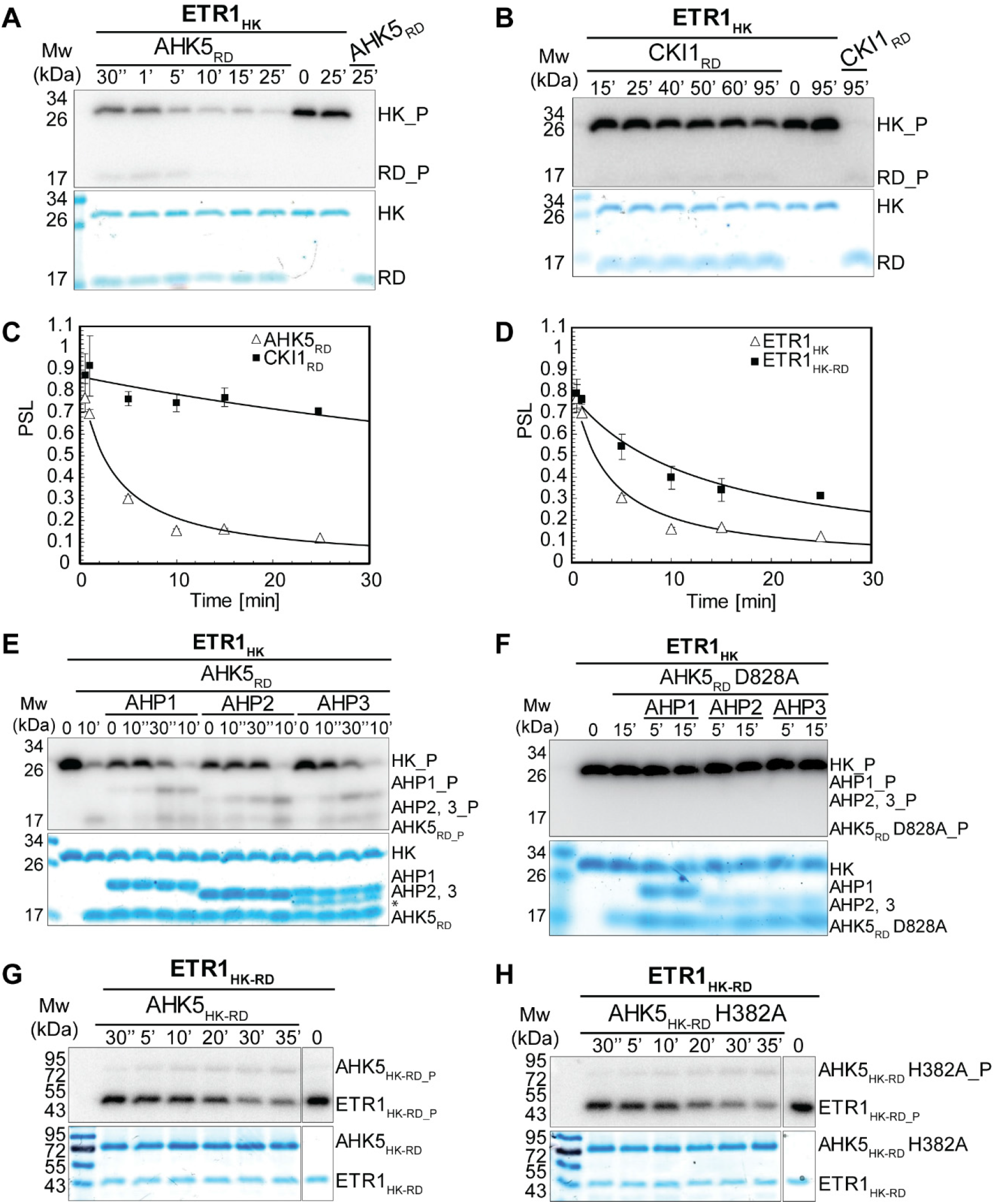
AHK5_RD_ is a specific substrate for the histidine kinase activity of ETR1 and AHK5 is able to transfer the phosphate from autophosphorylated ETR1_HK_ to AHPs independently of its own kinase activity. Phosphotransfer from ETR1_HK_ to AHK5_RD_ (**A**) and from ETR1_HK_ to CKI1_RD_ (**B**). Phosphotransfer is apparent after just 30 s of incubation in A. The last three lanes in A and B show ETR1_HK_ or AHK5_RD_/CKI1_RD_ only at the indicated time points after autophosphorylation. (**C**) Comparison of phosphotransfer kinetics from ETR1_HK_ to AHK5_RD_ (Δ) and CKI1_RD_ (■) estimated as GLM models based on the disappearance from ETR1_HK_ of the photostimulated luminescence (PSL) signal. Each point is the average of three replicates, error bars show standard error of the mean. (**D**) Comparison of phosphotransfer kinetics from ETR1_HK_ (Δ), and ETR1_HK-RD_ (■), respectively, to AHK5_RD_; curves estimated as in C. (**E**) Phosphate transfer from ETR1_HK_ to AHPs through AHK5_RD_ and **(F)** through AHK5_RD_ D828A. Time 0 means immediate addition of stopping buffer into the reaction. (**G**) Phosphotransfer from ETR1_HK-RD_ to AHK5_HK-RD_ and (**H**) AHK5_HK-RD_ H382A was performed as in A and B, respectively. In A-B and E-H, the top panels are autoradiograms following separation by SDS-PAGE. SDS-PAGE gels (12.5 %) stained with Coomassie-blue to show the amount of loaded proteins are shown below the autoradiograms. PageRuler Prestained Protein Ladder (ThermoFisher Scientific) was used as a molecular weight marker (Mw). The right-most lane in G and H shows autophosphorylated ETR1_HK-RD_ only. Asterisk (*) in E marks residual bacterial protein contamination.

To test for possible ETR1_HK_/AHK5_RD_/AHPs phosphorelay, equimolar amounts of AHPs and AHK5_RD_ were mixed and incubated with autophosphorylated ETR1_HK_. Phosphorelay manifested both via (i) disappearance of phosphorylated ETR1_HK_ and (ii) emergence of signal corresponding to phosphorylated AHK5_RD_ and the AHPs (Fig. 1E). Phosphotransfer from ETR1_HK_ to AHK5_RD_ and all AHPs was almost immediate (just after mixing together the individual reaction components, time “0” in Fig. 1E), confirming *in vitro* functionality and high efficiency of ETR1_HK_/AHK5_RD_/AHPs phosphorelay. Replacing the phosphoaccepting Asp 828 of AHK5_RD_ to the non phosphorylatable Ala (in the D828A mutant of AHK5_RD_) abolished phosphotransfer from ETR1_HK_ to both AHK5_RD_ and AHPs (Fig. 1F), thus confirming the His-to-Asp mechanism of the first transphosphorylation within the ETR1_HK_-induced phosphorelay. Importantly, the native AHK5 HK domain in the AHK5_HK-RD_ protein does not prevent the ability of ETR1_HK_ to phosphorylate AHK5_RD_ (Fig. 1G). AHK5_HK-RD_ is able to autophosphorylate and transfer the phosphate (via its own RD) to AHP2 (Supplemental Fig. 6). However, while the conserved His 382 of AHK5_HK_ is necessary for the autokinase activity of AHK5_HK-RD_ and AHK5-mediated phosphorylation of AHP2 (Supplemental Fig. 6), it is not required for the ETR1_HK_-mediated phosphorylation of AHK5_HK-RD_ (Fig. 1H). Altogether, the data convincingly demonstrate that ETR1HK is able to initiate the His-to-Asp-to-His phosphorelay, allowing the phosphorylation of AHPs via AHK5_RD_ independently of the HK activity of AHK5.

Despite the similarities in the overall structures, the ETR1_RD_ protein differs from other known bacterial, yeast and plant RDs in the orientation of its β3-α3 loop (also called γ-loop). Compared to all other (including bacterial and yeast) RDs of known structure, the γ-loop of ETR1_RD_ is flipped over to face the opposite side^8,9^ (Fig. 2A). Sequence analysis showed that the γ-loop of ETR1_RD_ differs from the γ-loop of AHK5_RD_ and other *Arabidopsis* RDs (Supplemental Fig. 3). To unravel the possible functional importance of this sequence variation, we introduced several mutations (namely G664V, V665L, E666D, and N667G) to rewire the sequence of the ETR1_RD_ γ-loop to resemble that of AHK5_RD_. Further, we mutated Val 711 of ETR1_RD_ to Phe, found in the corresponding position (903) of AHK5_RD_. F903 stabilizes the orientation of T884 during phosphorylation of AHK5_RD_ through F-T-coupling^10^. However, even the quintuple mutant ETR1_RD_ G664V V665L E666D N667G V711F was not able to accept phosphate from autophosphorylated ETR1_HK_ (Supplemental Fig. 7). To investigate the effect of introduced mutations on the structure of ETR1_RD_, we crystalized both wild type (WT) and mutant ETR1_RD_. The structure of WT ETR1_RD_ was determined at 1.5 Å resolution (PDB 7PN2). With the exception of the C-terminus position, only minor changes in the orientation of flexible β3-α4 loop were observed when compared to the previously published structure of ETR1_RD_ [solved at lower (2.5 Å) resolution, PDB 1DCF^9^ (Fig. 2A)]. In the quintuple mutant ETR1_RD_ G664V V665L E666D N667G V711F, the structure refined at 1.9 Å (PDB 7PN3) showed increased flexibility of the γ-loop when compared to the WT while retaining a similar orientation (Fig. 2B, Supplemental Table 1). To sum up, our structural data are in line with the functional assay, confirming the inability of ETR1_RD_ to accept phosphate from its native HK domain and suggest that molecular determinants of the observed structural and functional diversity are to be found outside the γ-loop of ETR1_RD_.

**Figure 2.**
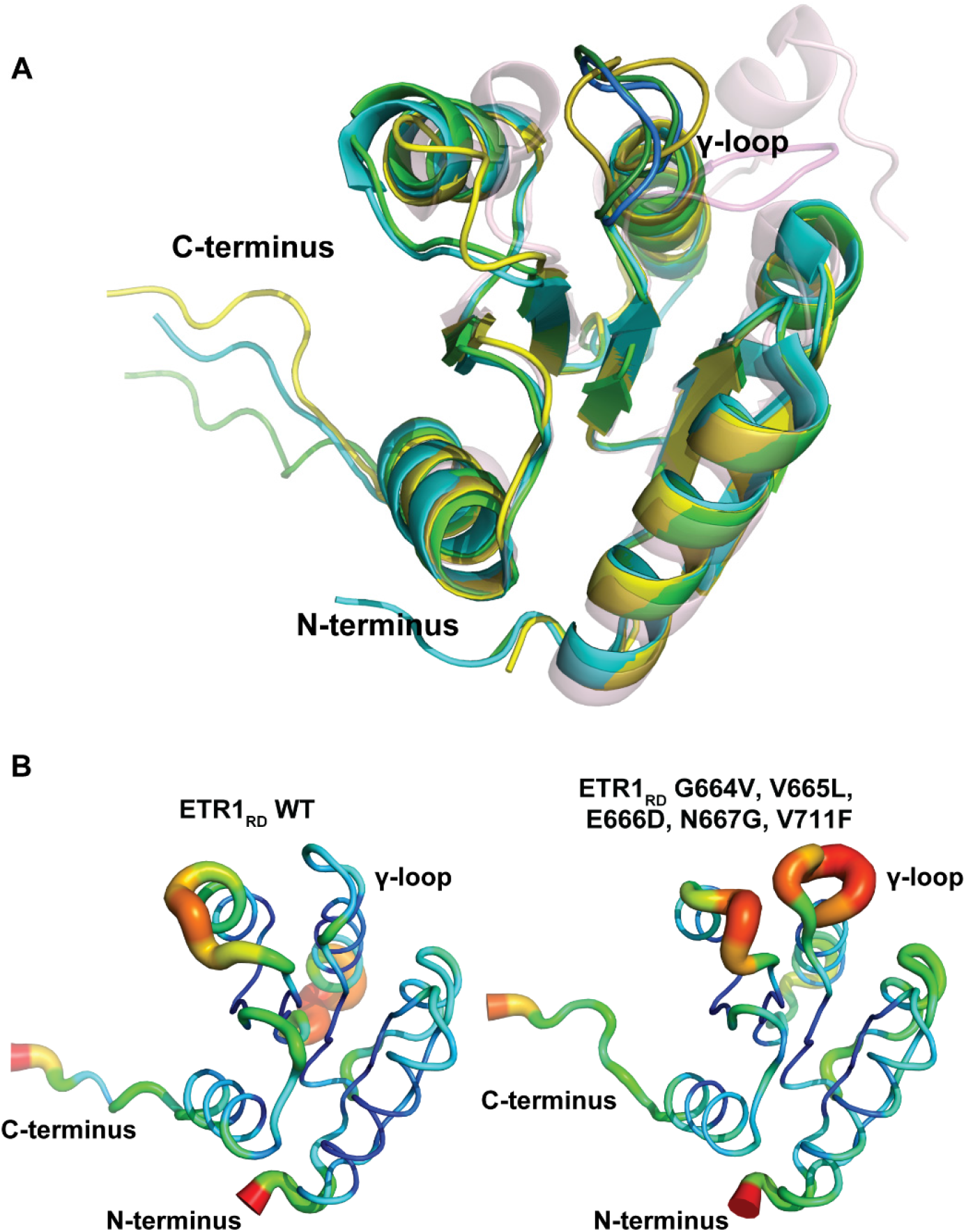
Mutant ETR1_RD_ with a β3-α3 loop rewired to resemble that of AHK5_RD_ is not able to accept phosphate from ETR1_HK_ due to structural differences. (**A**) Structural comparison of the ETR1_RD_, quintuple mutant ETR1_RD_ G664V, V665L, E666D, N667G, V711F and AHK5_RD_. Green – ETR1_RD_ WT (PDB 7PN2, this work), cyan – ETR1_RD_ (PDB 1DCF^9^), yellow – ETR1_RD_ G664V, V665L, E666D, N667G, V711F (PDB 7PN3, this work), light pink – AHK5_RD_ (PDB 4EUK^69^). Residues forming the γ-loop are shown in darker color. No significant changes were observed in the γ-loop position between WT and ETR1_RD_ G664V, V665L, E666D, N667G, V711F. (**B**) ETR1_RD_ (left) and ETR1_RD_ G664V, V665L, E666D, N667G, V711F (right) shown based on relative B-factor (blue for minimum and red for maximum). Increased flexibility of the β3-α3 loop of the mutant is highlighted using both color code and increased thickness of the line showing the peptide backbone.

Since ETR1_HK_ phosphorylates AHK5_RD_, we investigated the interaction between these proteins *in vitro* by MST and fluorescent anisotropy (FA) assays. Binding curves were measured by titration experiments where ETR1_HK_ or ETR1_HK-RD_ was added to fluorescently labelled AHK5_RD_. Results of both assays (Fig. 3A and Supplemental Fig. 8) show that ETR1_HK_ interacts with AHK5_RD_ albeit with low affinity. Interaction between the cytoplasmic part of ETR1 (ETR1 without transmembrane domain, ΔTM-ETR1) and full length AHK5 *in planta* has been assayed using FLIM-FRET. Statistically highly significant shortening (α = 9.05 x 10^-9^ and 5.5 x 10^-10^, respectively) of ΔTM-ETR1-GFP fluorescence lifetimes was detected in the presence of AHK5, tagged either N- or C-terminally with mRFP1 (Figs. 3B, C). Similar results were obtained in the FLIM-FRET assay using full length AHK5-GFP and ETR1-mCherry (Supplemental Fig. 9). These results prove that ETR1 associates with AHK5 *in planta*, allowing ETR1/AHK5/AHPs phosphorelay.

**Figure 3.**
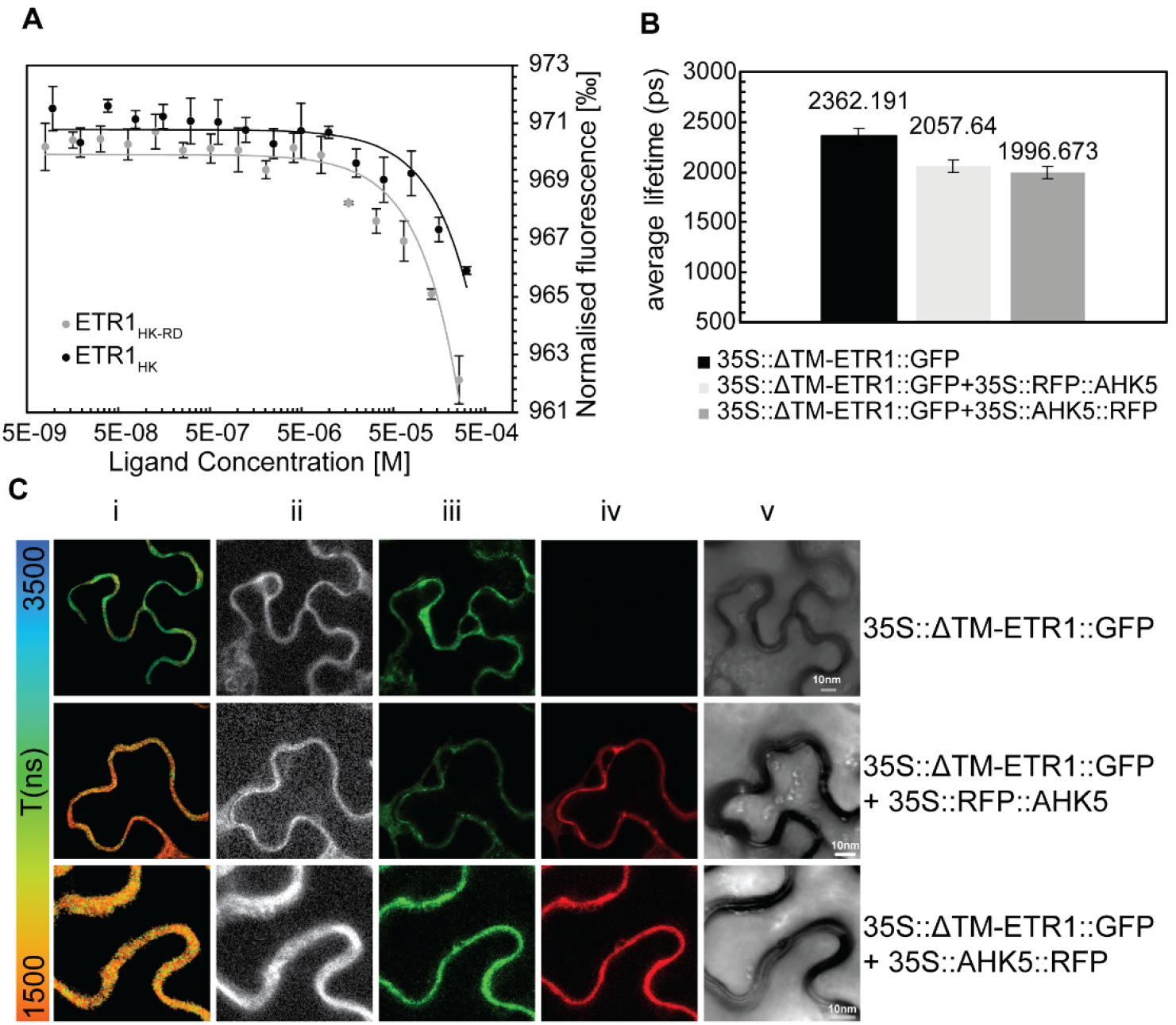
ETR1 interacts with AHK5 *in vitro* and *in vivo*. (**A**) Dose response curves for AHK5_RD_-Alexa Fluor 488 binding to ETR1_HK_ and ETR1_HK-RD_ using microscale thermophoresis (MST). The individual data points represent means of 3 replicates, error bars show standard deviation. Apparent K_D_ measured by MST for the interaction of AHK5_RD_ with ETR1_HK_ and ETR1_HK-RD_ is 166 ± 78 μM and 167 ± 50 μM, respectively. (**B**) Average fluorescence lifetime of ΔTM-ETR1::GFP measured in transiently transformed *N. benthamiana* leaves with and without AHK5 tagged at the N- or C-terminal with RFP. (**C**) Lifetime images and corresponding confocal images of ΔTM-ETR1::GFP transiently expressed in *N. benthamiana* leaves i) intensity weighted color-coded lifetime image of ΔTM-ETR1::GFP ii) photon intensity weighted lifetime image iii) GFP channel of the corresponding confocal micrograph iv) RFP channel of the corresponding confocal micrograph showing AHK5 expression v) transmitted light channel.

In order to assess the possible role of AHK5 in ethylene-regulated MSP signaling in *Arabidopsis*, we assayed the expression pattern of MSP reporter *TCSn:GFP*^11^ and its responsiveness to both cytokinins and ethylene in both *WT* and *ahk5-1* mutant backgrounds. As demonstrated previously^2,11^, the TCS signal locates mostly to the columella/lateral root cap (LRC) in mock-treated WT roots, while a weak signal is seen in the stele cells located proximally to the quiescent center (QC; Fig. 4A). Treatment with cytokinin ([5 μM 6-benzylaminopurine (BAP) for 24 h] upregulated the TCS signal throughout the root tip, i.e. in the stele, columella/LRC and epidermis/cortex of the root transition zone (TZ; Fig. 4A, Supplemental Figs. 10B-D). In line with a recent report^2^, 1-aminocyclopropane-1-carboxylic acid (ACC) – the rate-limiting precursor of ethylene biosynthesis – upregulated TCS preferentially in the root TZ. A relatively weaker activation (compared to mock-treated control) of MSP signaling was detected in the columella/LRC. In cases of combined treatment with cytokinin and ACC, most of the observed effects were cytokinin-like (Fig. 4A, Supplemental Fig. 10D). Strikingly, the absence of AHK5 almost completely abolished *TCSn:GFP* activity in the root, and only a weak signal was still apparent in the stele in mock-treated *ahk5-1* (Fig. 4B, Supplemental Fig. 10E). Further, the *TCSn:GFP* response of *ahk5-1* to cytokinin was comparable to WT, with an even slightly higher level of activation in the stele and root TZ. In contrast to this, ACC was able to induce TCS only in the root TZ and only weakly in the stele, but the signal was either weak or undetectable in the columella/LRC of *ahk5-1*, irrespective of whether ACC was present or not. This suggests that functional AHK5 is necessary for ethylene-induced MSP activity in the columella/LRC, but not for MSP induced by cytokinin in the same tissue. The role of ethylene in MSP signaling in the columella/LRC was further substantiated by downregulation of the *TCSn:GFP* signal in WT following treatment with either 2-aminoethoxyvinyl glycine (AVG), an inhibitor of ethylene biosynthesis or 1-methylcyclopropene (1-MCP), an ethylene antagonist that prevents ethylene binding to its receptors ETR1 and ERS1^12^ (Fig. 4C, Supplemental Figs. 10A, F).

**Figure 4.**
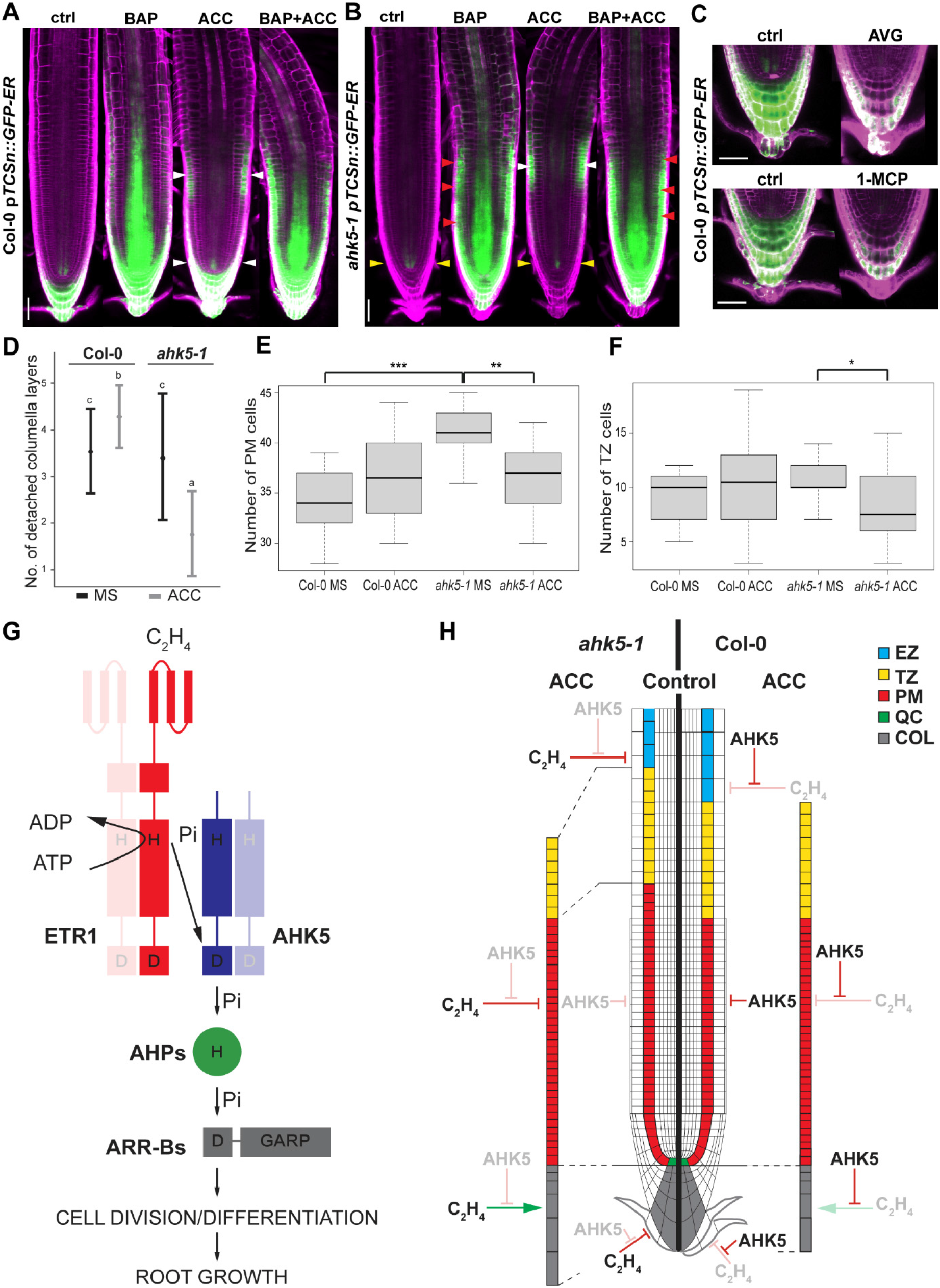
AHK5 is necessary for ethylene-mediated MSP activation in columella and lateral root cap. Expression pattern of ER-localized pTCSn-driven GFP in the WT Col-0 (**A, C**) and *ahk5-1* (**B**) backgrounds after (5 μM BAP, 5 μM ACC, 5 μM BAP + 5 μM ACC, 1 μM AVG, 2 μL·L^-1^ 1-MCP and 0.1 % DMSO/MS or air as control). The number of detached root cap layers of 10-day-old seedlings (**D**), proximal meristem size analysis (**E**) and number of TZ cells (**F**) were compared between Col-0 and *ahk5-1* seedlings grown on MS media ± 1 μM ACC. A model for the transphosphorylation crosstalk between ETR1 and AHK5 and its role in the hormonal regulation of root growth (**G** and **H**). The membrane signal from PI staining is shown in magenta; GFP is in green. White arrowheads in A and B mark the specific localization of the ACC-induced signal while the yellow arrowheads mark the positions where the ACC-induced signal is alleviated/missing in *ahk5-1* roots. Red arrowheads mark the stronger TCS signal in the epidermis/cortex upon BAP treatment. Scale bar represents 50 μm.

Ethylene was shown to control cell divisions in the quiescent center (QC) resulting in changes in the number of columella layers^13^. Accordingly, we found an additional columella cell layer in ACC-treated *ahk5-1* when compared to Col-0 (Supplemental Fig. 10G). Interaction between the activity of QC/columella initials, the number of root cap cells and their release as border cells in pea and border-like cells in *Arabidopsis*^14^ was shown to be under auxin and ethylene control^15,16^. We observed that while ACC slightly upregulated the number of border-like cells in WT, the opposite response, i.e. strong downregulation, was observed in ACC-treated *ahk5-1* (Fig. 4D).

Recently, ETR1-mediated MSP was shown to control root apical meristem (RAM) size by controlling cell differentiation in the root TZ^1,2^. Higher TCS induction by both cytokinin and ethylene observed particularly in the root TZ but also in the stele of *ahk5-1* (Supplemental Figs. 10D, E), implies a possible role for AHK5 also outside the columella/LRC. Accordingly, we found longitudinal expansion of the RAM proliferative region [also called proximal meristem (PM)^17^] in *ahk5-1* under control conditions. *ahk5-1* seedlings grown in the presence of ACC showed shortening of both RAM and TZ, which is in contrast to WT that was ACC-insensitive under these conditions (Figs. 4E, F). In agreement with these results, *ahk5-1* was more sensitive to ACC than WT also at the level of the root elongation (Supplemental Fig. 10H). Taken together, our data show that AHK5 is necessary for ethylene-mediated control of MSP signaling in the columella/LRC. Further our data underline that AHK5 is a negative regulator of cell number in the proliferative region of the RAM and attenuates both ethylene-regulated responses in columella/LRC as well as the ethylene-controlled RAM/TZ size.

## Discussion

Here we provide direct functional and structural evidence that although ETR1 is an active HK, ETR1_RD_ is unable to accept a phosphate moiety from phosphorylated ETR1_HK_. This could possibly be due to the high conformational rigidity and the different orientation of the ETR1_RD_ γ-loop^4,5^. Instead, ETR1 forms a complex with AHK5, another HK, and ETR1_HK_ phosphorylates AHK5_RD_ to initiate an ETR1/AHK5/AHPs phosphorelay. To our knowledge, this is the first example of signaling crosstalk mediated by transphosphorylation activity between two different sensor HKs (Fig. 4G) anywhere in the kingdoms employing MSP or related two-component signaling (bacteria, fungi and plants). A partially similar mechanism was described in *Pseudomonas aeruginosa*, where the RetS HK inhibits the activity of another HK GacS by three different mechanisms, each of which requires direct protein-protein interaction^18,19^. In contrast to the originally proposed model, the formation of a non-productive heterodimer between RetS and GacS has been recently excluded^20^. This type of interaction was recently described also in rice, where the ethylene sensor OsERS2 was shown to interact with the rice AHK5 orthologue MHZ1/OsHK1 and control its HK activity^21^. Nevertheless, transphosphorylation was not reported as a mechanism of the interaction either in the *Pseudomonas* or in the rice signaling systems, i. e., between RetS or OsERS2 and their downstream regulatory targets. Our data show direct interaction between ETR1 and AHK5, both *in vitro* and *in vivo*. Based on the known structural model of ETR1 and the complex between HK853 and its cognate RR RR468^22^, formation of a heterodimer between ETR1 and AHK5 seems to be the most plausible mechanistic model for explaining the observed phosphorelay from ETR1_HK_ to AHK5_RD_. While covalently-linked ETR1 homodimer formation is not necessary for ethylene binding^23^, further corroborating the possibility of functional heterodimer formation, the existence of higher order complexes (e.g. the dimer/dimer) cannot be excluded by our data (Fig. 4G).

ARR2, the type-B RR, was found to be phosphorylated in an ETR1-dependent manner^24^. Interestingly, AHK5 seems to act together with ETR1 in the H_2_O_2_-dependent control of stomata closure^25,26^ and ARR2 or AHP1, AHP2 and AHP3 were proposed to mediate the response downstream of ETR1 and AHK5, respectively^26,27^. AHP1-3 have been shown to physically interact with AHK5_RD_ *in planta*^10^ and with ARR2 in yeast^28,29^. Thus, our data showing the ability of ETR1_HK_ to phosphorylate AHP1, AHP2 and AHP3 through AHK5_RD_ are well in line with these observations, providing a mechanical window into the ETR1/AHK5/AHPs phosphorelay. Our results imply that AHK5 is a negative regulator of RAM size and a number of other ethylene-mediated responses including cell number regulation of columella/border-like cells and RAM and TZ shortening (Fig. 4H). This is also in agreement with the previously proposed negative role for AHK5 in ETR1-dependent ABA- and ethylene-mediated inhibition of root growth^30^. Apart from these, AHK5 has been shown to be a positive regulator of ethylene-induced stomata closure^25^. This implies that ETR1/AHK5 phosphorelay may act in a tissue-specific fashion, but the molecular mechanism behind this remains to be elucidated.

We demonstrate that AHK5 activity is critical for basal MSP signaling in the LRC/columella, representing most of the MSP signaling observed in the non-treated *Arabidopsis* roots. In LRC/columella cells, AHK5 mediates ethylene-induced, but not cytokinin-induced, MSP activation, and interferes with the ethylene-driven regulation of the number of columella cells and border-like cells detachment. Border and border-like cells are of paramount importance for root interaction with the soil microbiome and for protecting the root against biotic and abiotic stresses^31,32^. The release of border cells also facilitates root penetration through the soil by decreasing frictional resistance^33^. Recently, a role was proposed for ethylene in the inhibition of root growth through compacted soil^34^. The ethylene-inhibited release of border-like cells seems to represent another mechanism of ethylene-controlled root growth navigation in mechanically heterogeneous soil conditions, and as such, a potentially highly valuable crop breeding trait. However, our data show that AHK5 also acts outside the LRC/columella. Ethylene was shown to control root growth predominantly via the regulation of auxin biosynthesis and intercellular auxin distribution^35–38^. Importantly, MSP-regulated auxin accumulation specifically in the LRC was shown to be involved in the control of RAM size^39^. Thus, the AHK5-mediated regulation of auxin homeostasis in the LRC might be the mechanism through which AHK5 controls the RAM proliferation zone, and this could be partially ethylene-independent (Fig. 4G, H).

The absence of transmembrane and extracellular domains makes AHK5 an ideal partner for ETR1, which lacks a functional RD. This type of direct crosstalk between two sensor HKs seems to add a combinatorial aspect to the highly modular MSP signaling, thus widening the spectrum of signals that can potentially be integrated employing a single signaling mechanism. This type of signal integration seems to be important mainly for organisms living in highly variable conditions, including pathogenic bacteria as for instance the aforementioned RetS/GatS example, or typically for sessile organisms like plants. It still remains an open question as to whether AHK5 is able to integrate signal(s) from other HKs.

## Methods

### Phylogenetic analysis

Protein sequences of seven *Arabidopsis* histidine kinases (ETR1 [UniProtKB ID: P49333], AHK1 [Q9SXL4], AHK2 [Q9C5U2], AHK3 [Q9C5U1], AHK4 [Q9C5U0], AHK5 [Q3S4A7], and CKI1 [O22267]) were retrieved from the TAIR database (https://www.arabidopsis.org). C-terminal receiver domains were defined according to the SMART database^40^ and their sequence alignment was constructed using the Clustal Omega algorithm without additional constraints^41^. The phylogenetic tree was constructed using the Maximum Likelihood method based on the JTT matrix-based model in Mega 7 software^42^. Bootstrap values from 1,000 trials were used. The tree is drawn to scale, with branch lengths measured in the number of substitutions per site. All positions containing gaps and missing data were eliminated. There were a total of 115 positions in the final dataset.

### Cloning and mutagenesis

DNA constructs for protein expression of AHPs, AHK5_RD_, CKI1_RD_, AHK4_RD_, and ETR1_HK_ were prepared as reported earlier^8,43,44^. Two constructs were prepared for the production of ETR1_RD_ protein: (i) ETR1_RD_ in the expression vector pETm60-Ub2 (amino acid residues 609-738); the product was used in kinase assays and (ii) ETR1_RD_ in the pHYRSF53(Sumo)LA vector (amino acid residues 605-738); the product was used in kinase assays, for crystallization and in NMR experiments. The DNA construct in pETM60-Ub2 was prepared as reported^44^. The DNA fragments encoding ETR1_RD_ (605-738), ETR1_HK-RD_ (324-738), and AHK5_HK-RD_ (300-922) were amplified from cDNA by PCR using appropriate primers (see Supplemental Table 2) containing 5’-*Bam*HI and 3’-*Hind*III (for ETR1_RD_), 5’-*Nco*I and 3’-*Bam*HI (ETR1_HK-RD_), and 5’-*Bam*HI and 3’-*Eag*I (AHK5_HK-RD_) restriction sites. Amplified ETR1_RD_ and AHK5_HK-RD_ fragments were ligated into pHYRSF53(Sumo)LA and pFGRSF05^45^ (courtesy of Hideo Iwai, University of Helsinki) expression vectors, respectively and the ETR1_HK-RD_ fragment into pETM60-Ub2 vector^46^ (kindly provided by Vladimir V. Rogov, Goethe University Frankfurt). Expression plasmids were transformed into XL10-Gold Ultracompetent Cells (Agilent). DNA was extracted using ZR Plasmid Miniprep - Classic (Zymo Research) and resulting plasmids were verified by sequencing. All mutations listed in Supplemental Table 2 were introduced using the QuikChange Multi Site-Directed Mutagenesis Kit (Agilent) with appropriate mutagenic primers. AHK5_RD_ D828A and AHK5_HK-RD_ H383A were generated by one step mutagenesis. To generate the ETR1_RD_ quintuple mutant, three mutagenesis steps were implemented one after another: i) V711F, ii) G664V, V665L, and iii) E666D, N667G. DNA of ETR1_RD_ in pHYRSF53(Sumo)LA vector was used as a template in the first mutagenesis step. The mutated DNA from the previous step served as the template in each subsequent one. All introduced mutations were verified by sequencing.

### Protein expression and purification

The lengths and relative molecular weights of proteins used in this study are listed in Supplemental Table 3. AHP proteins were expressed and purified^43,44^ and ETR1_HK_ protein was obtained as previously described^8^. ETR1_RD_, ETR1_HK-RD_, and AHK5_HK-RD_ proteins were prepared as follows: Expression plasmids carrying ETR1_RD_ and ETR1_HK-RD_ genes were transformed into *E. coli* strain BL21(DE3)pLysS (Novagen, now Merck) and expression plasmid carrying AHK5_HK-RD_ gene into ER2566 (NEB) strain. Cells were cultured with shaking in Terrific broth (TB) medium at pH 7.5 for 3 h at 37 °C up to OD_600_ 0.6-0.8. Protein expression was induced using 1 mM isopropyl β-d-1-thiogalactopyranoside (IPTG) and continued for 5 h at 37 °C for ETR1_RD_ and overnight at 18 °C for ETR1_HK-RD_ and AHK5_HK-RD_ proteins. The cells were harvested by centrifugation at 3500 g for 20 min at 4 °C. The cell pellet was resuspended in lysis buffer (50 mM Tris-HCl, 0.5 M NaCl, 10 mM imidazole,10 % glycerol, 2.5 mM β-mercaptoethanol, 0.1 % Triton X-100, pH 7.9) and disrupted by sonication. After centrifugation at 47 000 g for 40 min at 4 °C, soluble proteins were purified by affinity chromatography using a HisTrap Ni-NTA column (ÄKTA pure FPLC system, GE Healthcare) equilibrated in 50 mM Tris-HCl, 0.5 M NaCl, 10 mM imidazole, 10 % glycerol, 2.5 mM β-mercaptoethanol, pH 7.5. Fusion proteins were eluted using a linear gradient of 0.5 M imidazole in equilibration buffer. Fusion tags were cleaved using S-TEV (Ub-His_6_) or UlpI (His_6_-TRX-SMT3) proteases overnight at room temperature while being dialyzed against TEV buffer (50 mM Tris-HCl, 0.5 mM EDTA, 1 mM DTT, pH 8) or at 4 °C while being dialyzed against PBS buffer, respectively. Proteins were purified from their cleaved fusion tags by a second affinity chromatography (using the same column and buffers), followed by size exclusion chromatography using a HiLoad 16/600 Superdex 75 prep-grade column equilibrated with 50 mM Tris-HCl, 150 mM NaCl, 1 mM EDTA, pH 7.5. Purified proteins were concentrated by ultrafiltration (Amicon Ultra system, Millipore) and stored at −80 °C. Mutated proteins (ETR1_RD_ quintuple mutant, AHK5_RD_ D828A and AHK5_HK-RD_ H382A) were overexpressed and purified in a manner identical to their wild-type versions. AHK5_RD_, CKI1_RD_, AHK4_RD_ and ETR1_RD_ (in pETM60-Ub2) were expressed according to a procedure published previously^44^ and purified as ETR1_HK-RD_ protein using lysis and purification buffers comprising 50 mM Na-phosphate instead of 50 mM Tris.

### Kinase assay

Prior to kinase assay all proteins were suspended in a kinase buffer (50 mM Tris-HCl, 10 % glycerol, 2 mM DTT, 5 mM MnCl_2_ and/or 5 mM MgCl_2_, pH 7.5) and the protein concentration was determined by the Bradford protein assay^47^ using bovine serum albumin (Serva) as a standard. The procedure for the kinase assay^48^ was adapted and carried out at room temperature (RT). Autophosphorylation was initiated by adding 1.2 μL [γ^32^P] ATP (10 μCi/mmol, M.G.P) to 120 μL of 3 μM protein and the mixture was incubated for 1 h at RT. Residual ATP was removed by desalting using Zeba™ Spin Desalting Columns, 7 K MWCO, 0.5 mL (ThermoFisher Scientific). Phosphorylated donor proteins were mixed with the phosphate acceptors in a 1:2 (3 μM:6 μM) molar ratio (or ratios of 1:4 and 1:6 for experiments presented in Supplemental Fig. 4). Total reaction volume was 10 μL, and the reactions were quenched at the times mentioned in the figures by adding 4×SDS-PAGE loading buffer: 0.2 M Tris-HCl, 0.4 M DTT, 277 mM, 8.0 % (w/v) SDS, 6 mM bromophenol blue, 4.3 M glycerol, supplemented with 80 mM EDTA. Phosphorylated proteins were resolved by SDS-PAGE (12.5 % gels or otherwise mentioned in the figure legend) without being boiled. Gels were wrapped in plastic foil and used for autoradiography by exposing to Fuji imaging plates. Incorporation of ^32^P was monitored using Typhoon FLA-7000 Scanner (GE Healthcare) and relative photostimulated luminescence (PSL) signals were quantified using MultiGauge software (FujiFilm). Phosphorylation depletion intensity was calculated by normalizing to the intensity of the band corresponding to autophosphorylated donor at time “0” alone (PSL_max_)^49^.

### Statistical analyses of phosphotransfer kinetics

The decline in the intensity of phosphorylated signals over time was approximated with an inverse model of the form *y* = 1/(*α+βx*) as it provided a better fit than a two-parameter exponential function. The comparison of the models for different proteins was performed via Generalized Linear Models with Gamma (GLM-g) error structure^50^.

### NMR spectroscopy

The ^15^N-labeled ETR1_RD_ samples were prepared according to a procedure published previously^51^, using M9 minimal medium supplemented with [15N]-NH4Cl. Cells (*E. coli* ER2566 carrying pHYRSF53(Sumo)LA vector with ETR1_RD_ gene) were harvested after overnight expression in 22 °C and purified as described above. Samples of 600 μL comprising 0.5 mM ^15^N-labeled ETR1_RD_ in 50 mM Na-phosphate-HCl buffer pH 7.5, containing 150 mM NaCl,1 mM DTT, 1 mM EDTA and 10 % deuterium oxide were used. Titrations with MgCl_2_ (steps of 0.25 mM, 0.5 mM, 0.75 mM, 1 mM, 1.25 mM, 1.5 mM, 1.75 mM, 2 mM, 3 mM, 4 mM, 5 mM, 8 mM, 10 mM, 30 mM, 50 mM) and MnCl_2_ (steps of 0.1 mM, 0.2 mM, 0.3 mM, 0.4 mM, 0.6 mM, 0.9 mM, 2 mM, 3 mM, 5 mM, 10 mM) were performed. Each titration step was measured as ^1^H-^15^N HSQC spectra. NMR spectra of MnCl_2_ titrations were acquired at 25 °C using a Bruker Avance III HD 600 MHz spectrometer, equipped with the cryogenic (^1^H-^31^P-^13^C-^15^N) inverse probe-head (Bruker) and MnCl_2_ titrations using a Bruker Avance III HD950 MHz spectrometer equipped with the cryogenic (^1^H-^13^C-^15^N) inverse probe-head.

### Microscale thermophoresis (MST) for protein-protein interaction analysis

To perform the MST experiment, purified AHK5_RD_ as a target was labelled with Alexa Fluor-488 (Thermo Fisher Scientific) according to the manufacturer’s instructions. The protein (20 mg·mL^-1^ in 50 mM HEPES, 300 mM NaCl, 5 % glycerol, pH 7.8) was mixed with 4 times the molar amount of Alexa Fluor-488 dye dissolved in 1/10 final volume of freshly prepared 1M NaHCO_3_ and stirred for 1 h at RT in darkness. The unbound dye was removed by ultrafiltration (Amicon Ultra system, Millipore) and the concentration of protein as well as the degree of labelling was checked. Fluorescently labelled AHK5_RD_ was kept constant at 92 nM in all MST measurements. ETR1_HK_ and ETR1_HK-RD_ solutions were prepared as a 2-fold dilution series of 16 concentrations (initial concentration of ETR1_HK_ was 315 μM and that of ETR1_HK-RD_ was 261 μM) in MST buffer containing 50 mM HEPES pH 7.8, 300 mM NaCl, 5 % glycerol and 0.015 % Fos-choline 16. Each reaction mixture was incubated for 10 min in a PCR-tube followed by thermophoresis in standard MST-grade glass capillaries in a Monolith NT.115 instrument (Nanotemper Technologies).

Measurements were performed under the blue fluorescence channel, with medium MST power, excitation power 80 % and constant 25 °C temperature. All results are averages of three independent repetitions.

### Fluorescent anisotropy assay

The assay was performed using a Fluoromax 4 spectrofluorometer (HORIBA, Jobin Yvon). Purified AHK5_RD_ protein was labeled with fluorescent Alexa-Fluor 484. The titration of 1 mM ETR1_HK_ solution into 25 nM AHK5_RD_-Alexa solution was performed in a 10 x 4mm quartz-glass cuvette with a chamber for a magnetic bar stirrer at 25°C. The interaction buffer was 50 mM HEPES, 200 mM NaCl, 5 mM MgCl_2_, 5 mM ATP, 0.05 % Tween, pH 7.5. All tests were repeated three times and averaged for each data point. The excitation wavelength was 492 nm and emission was monitored at 520 nm with a slit width of 5 nm. Integration time was 1 s. A cuvette was equilibrated with buffer containing labelled protein 10 min before each titration experiment by stirring (speed 5). Before each measurement point the sample was equilibrated in the cuvette with the ligand for 30 seconds. As a negative control the same measurement was performed adding buffer (as titrant) to AHK5_RD_ in the cuvette in the same volumes as ETR1_HK_ solution.

### *In planta* analyses

For *in planta* analyses, *Arabidopsis thaliana* wild-type (Col-0, N1092) and the *ahk5-1* mutant (N802391, SAIL_50_H11) were used. The seedlings were grown on vertically oriented plates with ½ strength Murashige and Skoog (MS) medium containing 1% sucrose in growth chambers (CLF Plant Climatics, GmbH). Six-day-old (6 DAG) seedlings were treated in liquid MS medium containing 5 μM benzylaminopurine (BAP, Duchefa), 5 μM 1-aminocyclopropane-1-carboxylic acid (ACC, Sigma), 5 μM BAP + 5 μM ACC, 1 μM aminoethoxyvinylglycine (AVG, Sigma) or 0.1% DMSO as control (mock) for 24 h prior to imaging and image processing for spatial-specific analyses of fluorescent reporter line *pTCSn::GFP-ER* (in Col-0^52^ and *ahk5-1* background) and columella cell count using ZEN blue (Carl Zeiss Microscopy, GmbH). 2 μL·L^-1^ 1-methylcyclopropene (1-MCP) was added to block ethylene signaling and the expression patterns of *pTCSn::GFP-ER*. 1-MCP was incorporated with air flow controlled by a gas mixing system (GMS; Photon System Instruments) inside closed chambers and air was used as control. For fluorescence visualization, roots were stained with PI and visualized using a ZEISS LSM780/880 confocal microscope. Quantification of reporter gene expression was described previously^52^ with slight modifications (see Fig. 4 and Supplemental Figure S10). At least 10 seedlings were evaluated in three independent experiments.

To monitor the number of detached root cap cell layers, five-day-old (5 DAG) seedlings were transferred to MS medium with or without 1 μM 1-aminocyclopropane-1-carboxylic acid, and grown on vertically oriented plates for five more days prior to imaging with the Axiolab 5 (Carl Zeiss Microscopy, GmbH). To determine the AHK5-dependent ethylene effect on the root cap, attached and detached columella, columella cell count analysis was done on *ahk5-1* and compared to that in Col-0 by counting the columella cell layers from the outermost root cap until the first differentiated columella cell layer and the detached root cap layers^13^. At least 15 seedlings were evaluated in three independent experiments.

The 6 DAG seedlings grown directly on +/- 1 μM ACC supplemented MS media was employed for root length, proximal meristem (PM) and transition zone (TZ) analysis. The root length was evaluated by tools of Figi. The size of PM and TZ was determined as a number of cortex cells filed from QC to the first elongated cell. Before imaging (LSM 800) the roots were fixed and stained with Schiff reagent and propidium iodide mixture. The length of the cortex cells was measured and evaluated via Cell-o-Tape plugin in Fiji^53^. The differences of cell length indicated borders of the PM, TZ and elongation zone. In statistics, changepoint detection is used to identify possible changes in a series of data. These changes may refer to mean, variance, trend, or other stochastic - statistical properties. In this work, an iterative Bayesian changepoint detection method is applied to identify significant changepoints in the mean root cells length as the cortex cells grow^54^. This method can be of great assistance in identifying root cell zones as it is effective when not too many changes occur and they are of a reasonable size^54^. The changepoint detection was implemented in R environment (v. 3.6.0) using the package “bcp”^55,56^.

Based on convention, all critical values are at α = 0.05. At least 15 seedlings were evaluated in three independent experiments.

### FLIM-FRET

All clones for FLIM-FRET experiments were constructed using Gateway™ technology (Invitrogen). The entry clones were obtained via BP-reaction in pDONR207. The binary vectors for expression of the GFP/RFP fusion proteins under the control of 35S promoter were constructed via LR-reaction using the corresponding entry clones and the destination vector pH7FWG2 for the ΔTM-ETR1 construct with C-terminal GFP, or pB7WGR2 for the AHK5 construct with N-terminal RFP and pB7RWG2 for the AHK5 construct with C-terminal RFP^57^. Primers used for cloning are in supplemental table S1. Construct preparation and tobacco infiltration procedures were identical to those described^58^. Gene silencing in *Nicotiana benthamiana* was suppressed by co-infiltrating the p19 protein from tomato bushy stunt virus cloned into pBIN61^59^.

FLIM-FRET data were acquired using a laser scanning confocal imaging microscope Zeiss LSM 780 AxioObserver equipped with an external In Tune laser (488-640 nm, < 3 nm pulse width, pulsed at 40 MHz, 1.5 mW), C-Apochromat 40 x water objective (NA 1.2) and the HPM-100-40 Hybrid Detector from Becker and Hickl GmbH. PMT detectors were used for acquiring confocal images using the same instrument. Photon counting was done using Simple-Tau 150N (Compact TCSPC system based on SPC-150N) with DCC-100 detector controller. GFP and RFP were excited at 490 nm wavelength from the In Tune laser and at 561 nm from DPSS-laser respectively, with appropriate laser power and PMT gain to capture confocal images of the donor and acceptor. Confocal images were processed in Zen 2.3 light version from Zeiss. After acquiring confocal images, the same cell was excited with only 490 nm In Tune laser with appropriate laser power to avoid any pile up effect, and the detector was changed to HPM100-40 to obtain FLIM data. Acquisition of FLIM data was done in SPCM 64 version 9.8 and SPCImage version 7.3 from Becker and Hickl GmbH was used for data analysis. To avoid background signal from chloroplasts, selected regions of interest (ROIs) containing signal in the corresponding confocal images were used for fluorescent lifetime calculation. A multiexponential decay model was used for fitting. In most cases two or three component fitting was applied to get the least χ^2^ parameter because of the background from the leaf specimen. Lifetime components with very low values below 500 ns were considered as background contribution and avoided for average lifetime calculations. FRET efficiencies were calculated using the equation

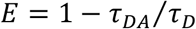

where E is energy transfer efficiency, *τ_DA_* is donor fluorescent lifetime in the presence of acceptor and *τ_D_* is the donor fluorescent lifetime in the absence of acceptor. Student’s t-test was used to evaluate the significant difference in the average lifetime between different groups.

### Protein crystallization

Initial crystallization conditions were obtained from the literature^51^. The conditions were further optimized by varying the temperature, the crystallization technique, and the presence of additives. For crystallization, the proteins were dissolved in 25 mM HEPES, 1 mM DTT, pH 7.5. Well-diffracting crystals were obtained using hanging drop vapor diffusion with 20 mg·lmL^-1^ ETR1_RD_ (15 mg·mL^-1^ for the ETR1_RD_ mutant) with 1.5 M Li_2_SO_4_, 0.1 M HEPES, pH 7.5 as a precipitant. The drops were prepared by mixing the protein and the precipitant in volumes of 1.5:1.5 μl, 1.5:3 μl and 3:1.5 μl. For the ETR1_RD_ WT, 0.1 M EDTA (0.5 μL) was added to each drop as an additive. The drops were equilibrated at 17 °C over a 1 ml reservoir of the precipitant. All crystals were cryo-protected in 30 % pentaerythritol ethoxylate and frozen in liquid nitrogen. Diffraction data were collected at 100 K at the BESSY II synchrotron (Helmholtz-Zentrum Berlin, Germany)^60^, beamlines 14.3 (crystals of wild-type ETR1_RD_) and 14.2 (ETR1_RD_ G664V, V665L, E666N, D667G, V711F).

### Structure determination

Diffraction images were processed using XDS^61^ or XDSAPP^62^ programs and converted to structure factors using the program Scala from package CCP4 v.7.0^63^ with 5 % of the data reserved for Rfree calculation. The structures of the complexes were solved by molecular replacement with Phaser^64^ (ETR1_RD_ G664V, V665L, E666D, N667G, V711F) or MOLREP^65^ (ETR1_RD_ WT), respectively. The ETR1_RD_ structure (PDB ID: 1DCF) was used as the initial coordinates for replacement. Refinement of the molecules was performed using REFMAC5^66^ alternated with manual model building in Coot v.0.8^66,67^. Water molecules were added by Coot and checked manually. The addition of alternative conformations, where necessary, resulted in final structures that were validated using the MolProbity^68^ (http://molprobity.biochem.duke.edu) validation server, and were deposited in the PDB database as 7PN2 and 7PN3 for the ETR1_RD_ WT and ETR1_RD_ G664V, V665L, E666D, N667G, V711F, respectively; molecular drawings were prepared using Pymol (Schrödinger, Inc.).

## Acknowledgements

We thank Prof. Stano Pekar (Masaryk University, Brno, Czech Republic) for statistical calculations. The work was supported by the Czech Science Foundation (19-24753S and 19-23108Y) and the Ministry of Education, Youth and Sports of the Czech Republic under the projects CZ.02.1.01/0.0/0.0/16_026/0008446, LTAUSA18161 and LTC19047. E.U. and E.Z. were supported by the Russian Science Foundation (20-14-00140). We acknowledge the Biomolecular Interactions and Crystallization Core Facility of CEITEC MU supported by the CIISB research infrastructure (LM2018127 funded by MEYS CR), the Proteomics Core Facility of CEITEC MU, the Plant Sciences Core Facility of CEITEC MU and the Josef Dadok National NMR Centre for their support in gathering the scientific data presented in this paper. We thank HZB for the allocation of synchrotron radiation beamtime. Part of this research was carried out at the light source PETRA III at DESY, a member of the Helmholtz Association (HGF). We acknowledge the core facility CELLIM of CEITEC supported by the Czech-BioImaging large RI project (LM2015062 funded by MEYS CR) for their support in gathering the scientific data presented in this paper.

## Author contributions

B.P. and Ja.H. conceived the research; A.S., A.R.C., B.P., M.Z., J.K., Z.J., A.J., M.H., and E.U. performed the research; A.S., A.R.C., B.P., M.Z., Jo.H., A.J., I.S., K.H., E.Z., L.Z., M.W., M.T., V.M. and Ja.H. analyzed the data; A.S., A.R.C., B.P., M.Z., A.J., E.Z., Jo.H., L.Z. and Ja.H. wrote the paper.

## Supplemental Material

Fig. 1. ETR1_RD_ is unable to mediate phosphotransfer in ETR1-mediated MSP signaling.

Fig. 2. Phylogenetic analysis of *Arabidopsis* RDs.

Fig. 3. Multiple sequence alignment of ETR1_RD_ and RDs from histidine kinases in *Arabidopsis thaliana*.

Fig. 4. ETR1_HK_ phosphorylates RDs of a different sensor HK.

Fig. 5. Kinetics of phosphotransfer from ETR1_HK_ to CKI1_RD_.

Fig. 6. Autophosphorylation of AHK5_HK-RD_ wild type (WT) and AHK5_HK-RD_ H382A.

Fig. 7. Mutant ETR1_RD_ with a β3-α3 loop rewired to resemble that of AHK5_RD_ is not able to accept phosphate from ETR1_HK_.

Fig. 8. ETR1_HK_ interacts with AHK5_RD_ *in vitro*.

Fig. 9. Full-length ETR1 and AHK5 interact *in vivo*.

Fig.10. AVG dilution series treatments with *pTCSn::GFP-ER*, representative ROI selection used for signal quantification and analysis of columella cell layers and root length.

Table S1. Data collection and refinement statistics for ETR1_RD_ proteins. Values in parentheses correspond to the highest resolution shell.

Table S2. Primer sequences used for cloning and mutagenesis. Restriction sites are underlined.

Table S3. List of proteins used.

