## Supplemental Data for "AHK5 mediates ETR1-initiated multistep phosphorelay in *Arabidopsis*"

### Supplemental Figures

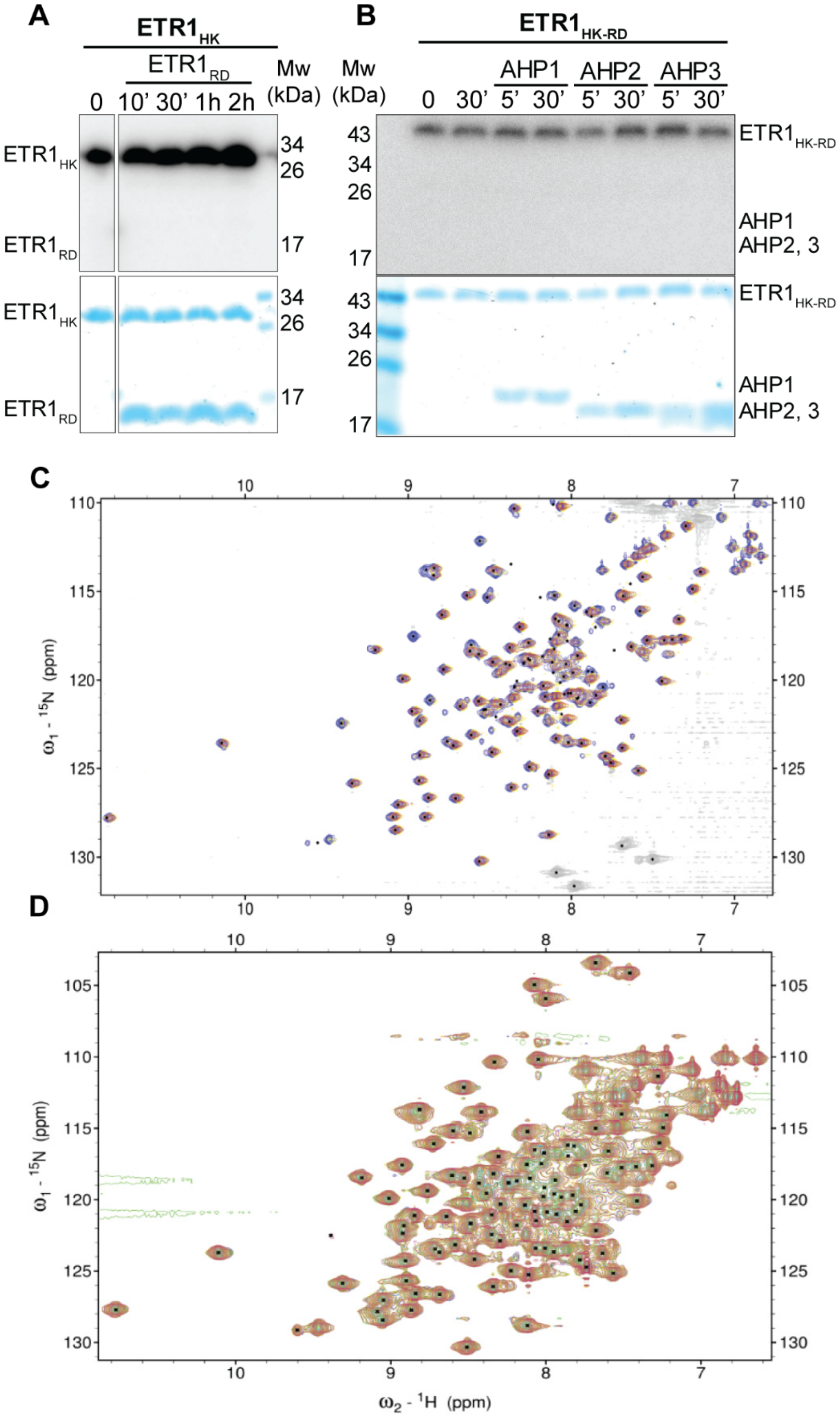


**Supplemental Figure 1. ETR1$_{RD}$ is unable to mediate phosphotransfer in ETR1-mediated MSP signaling.**

(**A**) Autophosphorylated $ETR1_{HK}$ was mixed with $ETR1_{RD}$ for the time periods indicated above the radiogram after which the reaction was terminated by addition of stop buffer. No transfer to $ETR1_{RD}$ was observed even after 2 hours. (**B**) Phosphotransfer assay between $ETR1_{HK\text{-}RD}$ and AHP1, 2 and 3, demonstrating inability to transfer phosphate from autophosphorylated $ETR1_{HK\text{-}RD}$ to AHPs after 5' and 30' of incubation. Top panels are autoradiograms following separation by SDS-PAGE. Coomassie-blue stained gels showing the amount of loaded proteins are shown below the radiograms. PageRuler Prestained Protein Ladder (ThermoFisher Scientific) was used as a molecular weight marker (Mw). (**C**) $^{1}H$-$^{15}N$ HSQC spectra of $MnCl_2$ titrations of $ETR1_{RD}$. Spectrum of free $ETR1_{RD}$ (blue), after addition of $MnCl_2$ to final concentration [0.4 mM (yellow)] and addition of $MnCl_2$ to final concentration [2 mM (red)]. The sample volume was 600 µL with a molarity of ~ 0.5 mM. (**D**) $^{1}H$-$^{15}N$ HSQC spectra of $MgCl_2$ titrations of $ETR1_{RD}$. Spectrum of free $ETR1_{RD}$ (blue), titrated with $MgCl_2$ at concentrations of 0.25 mM (orange), 0.5 mM (cyan), 0.75 mM (green), 1 mM (yellow), 2 mM (pink) and 5 mM (red). The sample volume was 600 µL with a molarity of 0.5 mM. Changes in peak position ($MgCl_2$) and peak intensity ($MnCl_2$) were very slight and did not show $ETR1_{RD}$ binding either $Mg^{2+}$ or $Mn^{2+}$ ions.

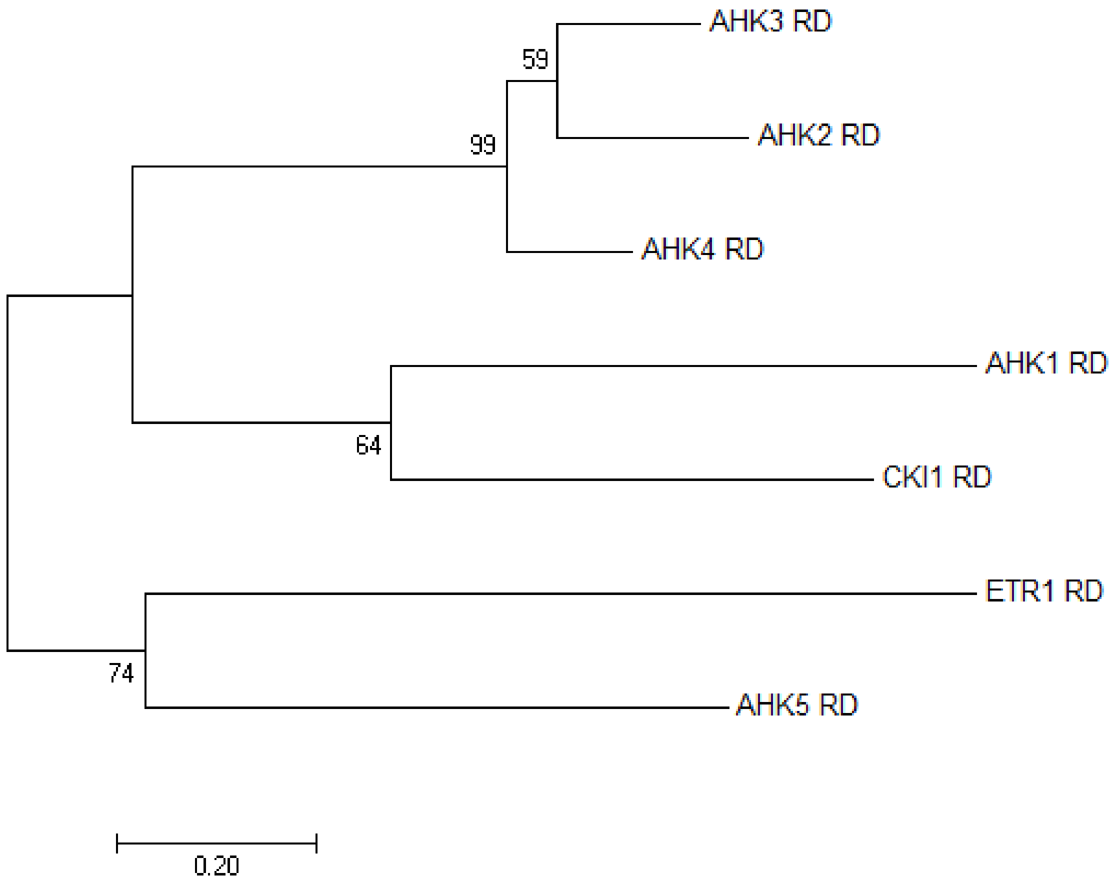


**Supplemental Figure 2. Phylogenetic analysis of *Arabidopsis* RDs.**

Phylogenetic tree was constructed using the Maximum Likelihood method and the JTT matrix-based model. Branch lengths correspond to the number of substitutions per site. All positions containing gaps and missing data were eliminated. There were a total of 115 positions in the final dataset. Evolutionary analyses were performed in MEGA7 software. Scale bar represents 0.2 substitutions per amino acid position.

```
ETR1_RD 610   LKVLVMDENGVSRMVTKGLLVHLGCEVT-TVSSNEECLRVVSH-----------------   652
AHK5_RD 778   PKILLVEDNKINIMVAKSMMKQLGHTMD-IANNGVEAITAI-------------------   818
AHK1_RD 986   IRILLAEDTPVLQRVATIMLEKMGATVTAVWD-GQQAVDSLNYKSINAQAPTEEHKSFEE   1045
CKI1_RD 1044  KRVLVVDDNFISRKVATGKLKKMGVSEVEQCDSGKEALRLVT-EGL--------------   1089
AHK2_RD 1035  KQILVVDDNLVNRRVAEGALKKYGAIVT-CVESGKAALAMLK------------------   1076
AHK3_RD 890   RKILIVDDNNVNLRVAAGALKKYGADVV-CAESGIKAISLLK------------------   931
AHK4_RD 945   KKILVVDDNIVNRRVAAGALKKFGAEVV-CAESGQVALGLLQ------------------   986
               :: : :       *:   : : *       . .  .:  :
                                       γ-loop
ETR1_RD 653   -----------------EHKVVFMDVCMPGVENYQIALRIHEKFTKQ-------------   683
AHK5_RD 819   --------------NSSSYDLVLMDVCMPVLDGLKATRLIRSYEETGNWNAAIEAGVDIS   865
AHK1_RD 1046  ETANKVTTRETSLRNSSPYDLILMDCQMPKMDGYEATKAIRRAEIGT-------------   1093
CKI1_RD 1090  -----TQREEQGSVDKLPFDYIFMDCQMPEMDGYEATREIRKVEKSYGVRT---------   1136
AHK2_RD 1077  --------------PPHNFDACFMDLQMPEMDGFEATRRVRELEREINKKIA--SG----   1117
AHK3_RD 932   --------------PPHEFDACFMDIQMPEMDGFEATRRIRDMEEEMNKRIK--NG----   972
AHK4_RD 987   --------------IPHTFDACFMDIQMPQMDGFEATRQIRMMEKETKE-----------   1022
                                .   .:*  *  ::. : :  ::

ETR1_RD 684   -----------RHQRPLLVALSGNT-DKSTKEKCMSFGLDGVLLKPVSLDNIRDVL----   725
AHK5_RD 866   TSENEQVCMRPTNRLP-IIAMTANT-LAESSEECYANGMDSFISKPVTLQKLRECL----   917
AHK1_RD 1094  -----------ELHIP-IVALTAHA-MSSDEAKCLEVGMDAYLTKPIDRKLMVSTI----   1114
CKI1_RD 1137  ---------------P-IIAVSGHDPGSEEARETIQAGMDAFLDKSLNQLANVI------   1192
AHK2_RD 1118  EVSAEMFCKFSSWHVP-ILAMTA-DVIQATHEECMKCGMDGYVSKPFEEEVLYTAV----   1169
AHK3_RD 973   EALIVENGNKTSWHLP-VLAMTA-DVIQATHEECLKCGMDGYVSKPFEAEQLYREV----   1024
AHK4_RD 1023  -------KTNLEWHLP-ILAMTA-DVIHATYEECLKSGMDGYVSKPFEEENLYKSV----   1067
                             * ::* :          .    *::. : * .
```

**Supplemental Figure 3. Multiple sequence alignment of $ETR1_{RD}$ and RDs from histidine kinases in *Arabidopsis thaliana*.**

Proteins used for alignments are: ETR1 [P49333], AHK1 [Q9SXL4], AHK2 [Q9C5U2], AHK3 [Q9C5U1], AHK4 [Q9C5U0], AHK5 [Q3S4A7], CKI1 [O22267] UniProt accession numbers are in brackets. Amino acids marked in red were subject to mutagenesis for rewiring $ETR1_{RD}$ into $AHK5_{RD}$. The conserved aspartate is marked in green. The position of the γ-loop is highlighted in bold. The alignment was generated using Clustal Omega software with manual adjustment.

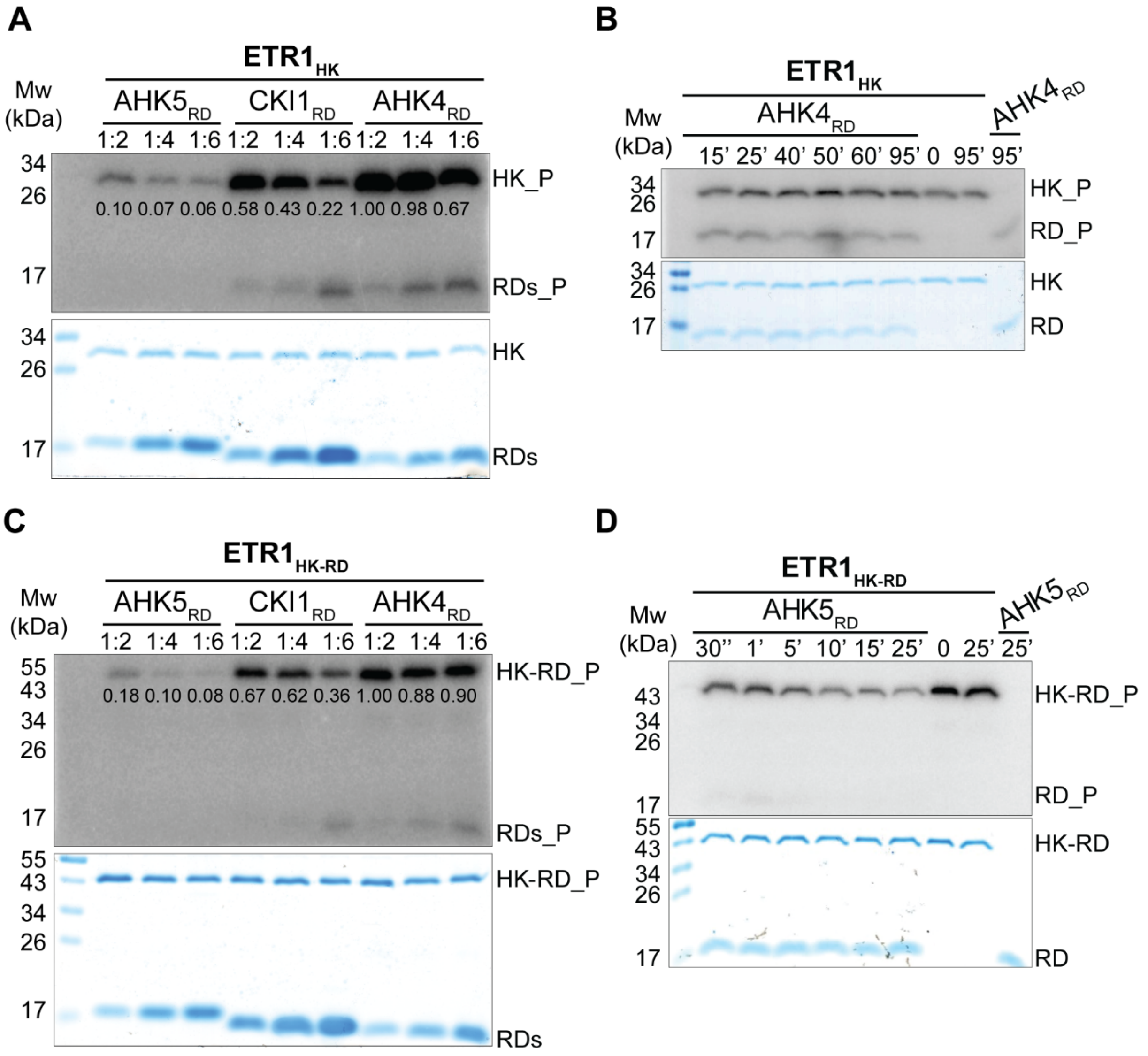


**Supplemental Figure 4. $ETR1_{HK}$ phosphorylates RDs of a different sensor HK.**

(**A**) Phosphotransfer between phosphorylated $ETR1_{HK}$ and three RDs in molar ratios of 1:2 (3 μM:6 μM), 1:4 (3 μM:12 μM) and 1:6 (3 μM:18 μM) after 30 minutes. The numbers on the radiograms indicate the signal of autophosphorylated $ETR1_{HK}$ remaining after addition of RDs normalized to the highest photostimulated luminescence signal in each radiogram ($ETR1_{HK}$ in the presence of $AHK4_{RD}$ at molar ratio 1:2). Phosphate transfer is shown by a decrease of $ETR1_{HK}$ signal and/or appearance of the phosphate acceptor signal ($AHK5_{RD}$, $CKI1_{RD}$ or $AHK4_{RD}$). The most intense phosphotransfer (the highest loss of $ETR1_{HK}$ phosphorylation corresponding to ≤ 10% of the strongest signal) is seen with $AHK5_{RD}$, then to $CKI1_{RD}$ and $AHK4_{RD}$. Higher molar ratio between phosphate donor and acceptor leads to more apparent phosphate transfer. (**B**) Phosphotransfer from $ETR1_{HK}$ to $AHK4_{RD}$ in a molar ratio 1:2. (**C**) Phosphotransfer between autophosphorylated $ETR1_{HK\text{-}RD}$ and individual RDs, same as in A. (**D**) Phosphotransfer between autophosphorylated $ETR1_{HK\text{-}RD}$ and $AHK5_{RD}$ in a molar ratio 1:2. The last three lanes (B and D) show $ETR1_{HK}$/$ETR1_{HK\text{-}RD}$ or $AHK5_{RD}$/$AHK4_{RD}$ only at the indicated time points after autophosphorylation. Top panels show autoradiograms following separation by gradient SDS-PAGE (4–20 % Mini-PROTEAN® TGX™ Precast Protein Gels). Coomassie-blue stained gels showing the amount of loaded proteins are shown below the radiograms. PageRuler Prestained Protein Ladder (ThermoFisher Scientific) was used as a molecular weight marker (Mw).

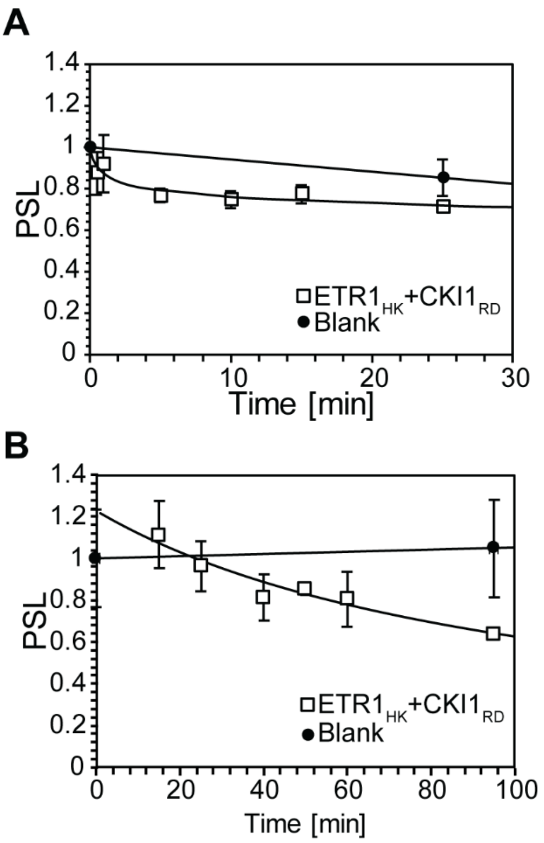


**Supplemental Figure 5. Kinetics of phosphotransfer from $ETR1_{HK}$ to $CKI1_{RD}$.**

$ETR1_{HK}$ phosphorylation status quantified using photostimulated luminescence (PSL) measured up to 25 min (**A**) and 95 min (**B**). Phosphorylation levels of $ETR1_{HK}$ were normalized to blank (autophosphorylated $ETR1_{HK}$ only) at T0. Only negligible phosphotransfer from $ETR1_{HK}$ to $CKI1_{RD}$ was detectable, particularly after 25' of incubation when compared with phosphotransfer from $ETR1_{HK}$ to $AHK5_{RD}$ (see Fig. 1D, F, main text). Each point is the average from three experiments; error bars show standard error of the mean.

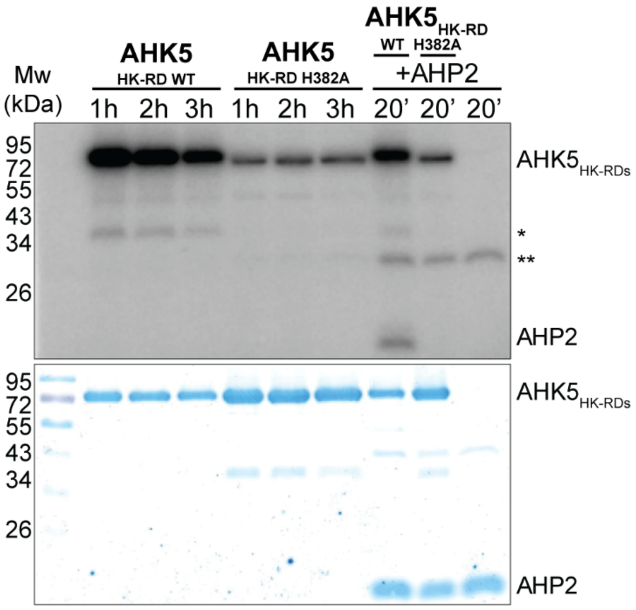


**Supplemental Figure 6. Autophosphorylation of AHK5$_{HK-RD}$ wild type (WT) and AHK5$_{HK-RD}$ H382A.**

Autophosphorylation assay using purified AHK5$_{HK-RD}$ WT as well as AHK5$_{HK-RD}$ H382A performed for up to 3 h. Compared to WT, AHK5$_{HK-RD}$ H382A shows only residual autokinase activity (possibly due to non-productive binding of [γ$^{32}$P] ATP). This is not sufficient to trigger phosphorelay to AHP2 (see the last three lanes). As a negative control, AHP2 was incubated with [γ$^{32}$P] ATP only, the transphosphorylation assay was stopped after 20 min. Asterisks mark residual bacterial proteins after purification.

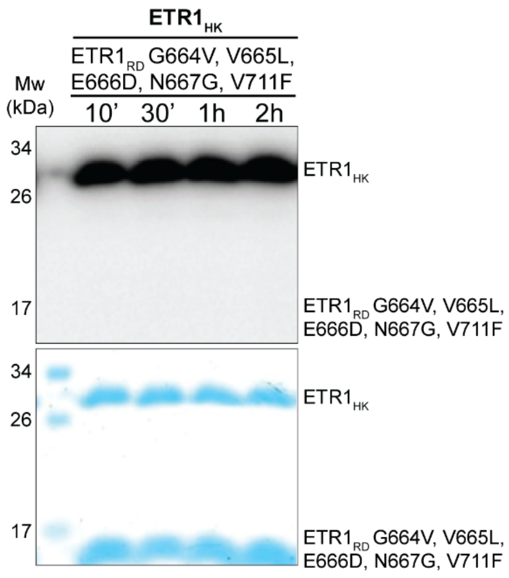


**Supplemental Figure 7. Mutant ETR1$_{RD}$ with a β3-α3 loop rewired to resemble that of AHK5$_{RD}$ is not able to accept phosphate from ETR1$_{HK}$.**

ETR1$_{HK}$ autophosphorylated for 1h, desalted to remove residual ATP, was mixed with the ETR1$_{RD}$ G664V, V665L, E666D, N667G, V711F quintuple mutant for the durations indicated above. The top panel shows the autoradiogram following separation by 12.5 % SDS-PAGE, Coomassie-blue stained gel is shown below the radiogram. No transfer to ETR1$_{RD}$ mutant was observed even after 2 hours of incubation.

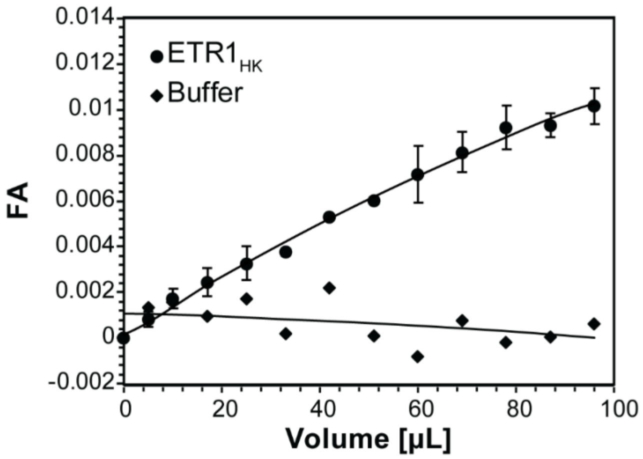


**Supplemental Figure 8. $ETR1_{HK}$ interacts with $AHK5_{RD}$ *in vitro*.**

Fluorescence anisotropy (FA) of Alexa 488-labeled $AHK5_{RD}$ mixed with increasing amounts of $ETR1_{HK}$, both at 25 nM concentration (●) and identical volumes of reaction buffer as a negative control (♦). The almost linear trend of binding isotherms suggests very weak interaction between the proteins. The equilibrium dissociation constant could not be determined as we were not able to reach saturation due to limited solubility of $ETR1_{HK}$. Error bars show standard error of mean calculated from three replicates.

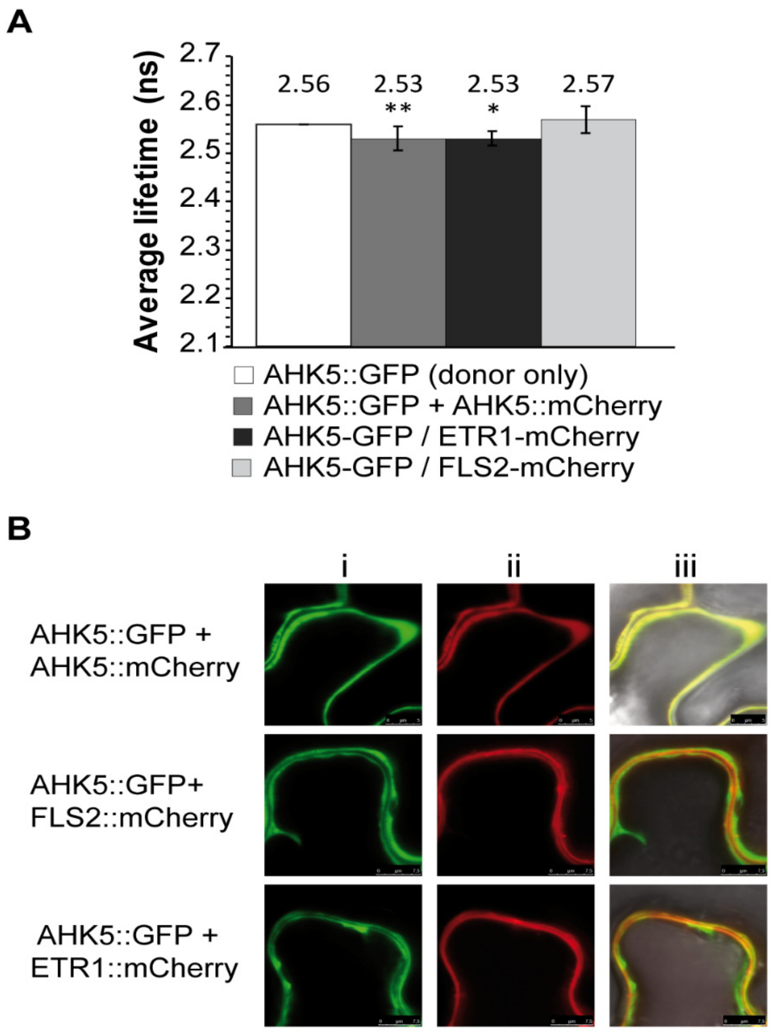


**Supplemental Fig. 9. Full-length ETR1 and AHK5 interact *in vivo.***

(**A**) Average fluorescence lifetime of AHK5::GFP measured in transiently transformed *N. benthamiana* leaves with mCherry labelled ETR1, AHK5 (positive control) or FLS2 (negative control), t-test with AHK5-GFP donor only ***P < 0.001, **P < 0.01, *P < 0.05. (**B**) Lifetime images and corresponding confocal images of AHK5::GFP transiently expressed in *N. benthamiana* leaves along with labelled ETR1, AHK5 or FLS2 i) GFP channel of the confocal micrograph ii) mCherry channel of the corresponding confocal micrograph showing AHK5, FLS2 or ETR1 expression iii) overlay of GFP, mCherry and transmitted light channel.

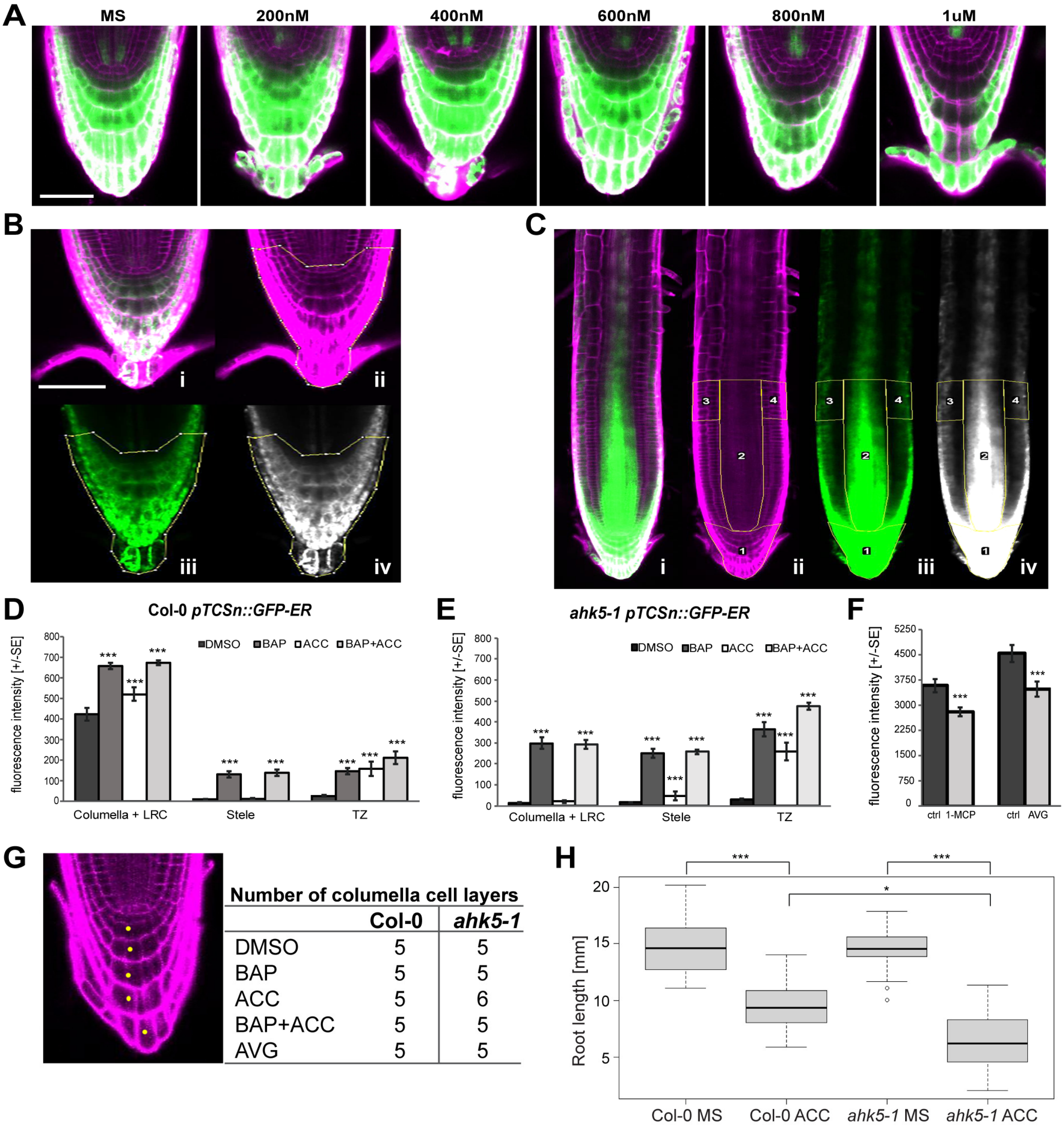


**Supplemental Figure 10. AVG dilution series treatments with *pTCSn::GFP-ER*, representative ROI selection used for signal quantification and analysis of columella cell layers and root length.**

(**A**) Six-day-old *pTCSn::GFP-ER* seedlings after 24 h treatment with increasing concentrations of AVG. (**B, C**) Representative images of *pTCSn::GFP-ER* roots showing the ROI selected for GFP signal quantification. In B and C, (i) GFP merged with propidium iodide (PI) stained cell walls, (ii) PI and (iii) GFP channels, (iv) denoised image of GFP signal in grayscale done in Fiji. These ROI were applied to the denoised GFP channel and the multi-measure tool was used for quantification of grey values. (**B**) ROI selection for columella and lateral root cap (LRC) done on DMSO-treated root. (**C**) ROI selection for columella and LRC (1), the stele (2), and the left (3) and right (4) transition zone indicated on a processed image of BAP treated root. Only the PI channel was used for ROI selection. GFP signal is in green, PI is in magenta. Scale bars correspond to 50 µm. (**D**-**F**) Relative fluorescence intensity quantification data from Col-0 *pTCSn::GFP* (**D, F**) and *ahk5-1 pTCSn::GFP* (**E**) expression patterns after hormone treatments. Statistical significance (t-test) differences between control and treated roots at alpha < 0.001 is denoted by asterisks (***); error bars show standard error (D-F). (**G**) Number of columella cell layers in 10-days old seedlings. (**H**) Root length of six-day-old seedlings grown on MS with or without 1 µM ACC. The Dunn-Kruskal-Wallis multiple comparison of p-values adjusted with the Holm method is denoted by asterisks.

**Table S1. Data collection and refinement statistics for ETR1$_{RD}$ proteins.** Values in parentheses correspond to the highest resolution shell.

| **Molecule** | **ETR1$_{RD}$** | **ETR1$_{RD}$ G664V V665L E666D N667G V711F** |
|---|---|---|
| PDB ID | 7PN2 | 7PN3 |
| *Data collection* | | |
| Beamline | BESSY II 14.3 | BESSY II 14.2 |
| Wavelength (Å) | 0.8950 | 0.9184 |
| Space group | $P4_32_12$ | $P4_32_12$ |
| Unit-cell parameters | | |
| a and b (Å) | 50.75 | 43.98 |
| c (Å) | 111.92 | 110.37 |
| α = β = γ [°] | 90 | 90 |
| Resolution range (Å) | 46.22-1.47 (1.55-1.47) | 43.98-1.92 (2.02-1.92) |
| Reflections measured | 391002 (39948) | 123444 (17917) |
| Unique reflections | 25479 (3274) | 8908 (1244) |
| Completeness (%) | 98.3 (89.6) | 100.0 (100.0) |
| $CC_{1/2}$ | 100.0 (77.9) | 99.8 (63.6) |
| $R_{merge}$† | 0.040 (1.730) | 0.179 (1.894) |
| <I/σ (I) > | 31.1 (1.5) | 10.1 (1.5) |
| Multiplicity | 15.3 (12.2) | 13.9 (14.4) |
| Wilson B factor (Å$^{-2}$) | 23.4 | 21.3 |
| *Refinement* | | |
| Reflections used | 24178 (1492) | 8419 (598) |
| Reflections used for $R_{free}$ | 1232 (62) | 436 (31) |
| R factor‡ (%) | 21.4 (35.0) | 20.4 (31.0) |
| $R_{free}$‡ (%) | 26.0 (32.3) | 26.8 (29.5) |
| R.m.s.d. bond lengths (Å) | 0.009 | 0.011 |
| R.m.s.d. bond angles (°) | 1.493 | 1.676 |
| R.m.s.d. chirals (Å$^3$) | 0.078 | 0.075 |
| No. of non-H atoms (total) | 1218 | 1090 |
| No. of water molecules | 106 | 44 |
| No. of ions | 4 | 0 |
| Mean B factor (Å$^2$) | | |
| Non-H atoms | 32.1 | 34.8 |
| Water molecules | 38.7 | 40.0 |
| Ions | 52.6 | - |
| Ramachandran plot (%) | | |
| Residues in most favourable regions (%) | 100.0 | 98.5 |
| Residues in allowed regions (%) | 0.0 | 1.5 |
| Rotamer outliers (%) | 0.0 | 2.5 |

† $R_{merge} = \sum|I_i - \langle I \rangle|/\sum I_i$, where $I_i$ is the intensity of observation and <I> is the mean value for that reflection.

‡ R factor $=\sum||F_o(h)| - |F_c(h)|| / \sum_h F_o(h)$, where $F_o$ and $F_c$ are the observed and calculated structure-factor amplitudes, respectively.

**Table S2. Primer sequences used for cloning and mutagenesis. Restriction sites are underlined.**

| | |
|---|---|
| **Cloning** | |
| **Gene** | **Primer sequences** |
| **ETR1$_{RD}$** (609-738) | Fw: 5´ GCTACCCATGGAAGGACTTAAGGTT-3`<br>Rev: 5´ CTGGGATCCTATTACATGCCCTC-3` |
| **ETR1$_{RD}$** (605-738) | Fw: 5’-CAGGATCCAATTTCACTGGACTTAAGGTTC-3’<br>Rev: 5’-TTAAAGCTTACATGCCCTCGTACAGTACC-3’ |
| **ETR1$_{HK}$** | Fw: 5’-TAGGATCCATGGAGCAGAATGTTGCTCTTG-3’<br>Rev: 5’-AGTAAAGCTTAATGTCGGGGAATGGCTGGAAC-3’ |
| **ETR1$_{HK-RD}$** | Fw: 5’-GGACCCATGGAACAGAATGTTG-3’<br>Rev: 5’-GATGGATCCTTACATGCCCTCGTA-3’ |
| **AHK5$_{HK-RD}$** | Fw: 5’-GTAGTGGATCCATGGAATCAGAACTGAAC-3’<br>Rev: 5’-GATCGGCCGTCAGTGCAAATACTGTTG-3’ |
| **AHK5$_{RD}$** | Fw: 5’-GGCCATGGAAACATCAAAGCCAA-3’<br>Rev: 5’-CGCGGATCCTAGTGCAAATACTGT-3’ |
| **CKI1$_{RD}$** | Fw: 5’-GGGCCATGGAAGGAAAGAGAGTTCTTGTG-3’<br>Rev: 5’-AGCGGATCCTAGTGACGTTTGCTTTCGATT-3’ |
| **AHK4$_{RD}$** | Fw: 5’-GGGCCATGGAAGGGAAGAAGATTCTTG-3’<br>Rev: 5’-TGCGGATCCTACGACGAAGGTGAGATA-3’ |
| **AHP1** | Fw: 5’-CCGGAATTCATGGATTTGGTTCAGAAGCAG-3’<br>Rev: 5’-CCGGAATTCTCAAAATCCGAGTTCGACGGC-3’ |
| **AHP2** | Fw: 5’-CCGGAATTCATGGACGCTCTCATTGCTCAG-3’<br>Rev: 5’-CCGGAATTCTTAGTTAATATCCACTTGAGG-3’ |
| **AHP3** | Fw: 5’-CCGGAATTCATGGACACACTCATTGCTCAG-3’<br>Rev: 5’-CCGGAATTCTTATATATCCACTTGAGGGAT-3’ |
| **Mutagenesis** | |
| **AHK5$_{HK-RD}$ H382A** | Fw: 5’-GATGCTAGCGACGATGTCTGCGGAGATAAGGTCACCATTG-3’<br>Rev: 5’-CAATGGTGACCTTATCTCCGCAGACATCGTCGCTAGCATC-3’ |
| **AHK5$_{RD}$ D828A** | Fw: 5’-ACGATCTGGTACTCATGGCGGTGTGCATGCCGGTGCTC-3’<br>Rev: 5’-GAGCACCGGCATGCACACCGCCATGAGTACCAGATCGT-3’ |
| **ETR1$_{RD}$ V711F** | 5’-GAGCTTTGGTCTAGACGGTTTTTTGCTCAAACCCGTATCAC-3’ |
| **ETR1$_{RD}$ G664V V665L** | 5’-GGACGTGTGCATGCCCGTGCTGGATGGCTACCAAATCGCTCTCC-3’ |
| **ETR1$_{RD}$ E666D N667G** | 5’-CGTGTGCATGCCCGGGGTCGATGGCTACCAAATCGCTCTCCGTATT-3’ |
| **Gateway Cloning** | |
| **5'-attB1-ETR1ΔTM** | 5’-AAAAAGCAGGCTTAATGGAGGGTAACATTTGGATTGA-3’ |
| **5'-attB2-ETR1ΔTM** | 5’-AGAAAGCTGGGTGCATGCCCTCGTACAGTACC-3’ |
| **5'-attB1-AHK5** | 5’-AAAAAGCAGGCTTAATGGAGTCAGAGCTCACAGT-3’ |

| | |
|---|---|
| **5'-attB2-AHK5** | 5’-AGAAAGCTGGGTGGTGCAAATACTGTTGCAAACAC-3’ |
| **5'-attB2-AHK5 with STOP codon** | 5’-AGAAAGCTGGGTTCAGTGCAAATACTGTTGCAAACAC-3’ |

**Table S3. List of proteins used.**

| Protein name | Length (start – end), Mw | References |
|---|---|---|
| **$ETR1_{RD}$** | (609-738), 14953 Da | Borkovcová et al., Phytochemistry 100 (2014) 6–15 |
| **$ETR1_{RD}$** | (605-738), 15014 Da | - |
| **$ETR1_{HK}$** | (324-604), 30815 Da | Otrusinova et al., J. Biol. Chem. (2017) 292(42) 17525–17540 |
| **$ETR1_{HK\text{-}RD}$** | (324-738), 45940 Da | - |
| **$AHK5_{HK\text{-}RD}$** | (300-922), 62967 Da | - |
| **$AHK5_{HK\text{-}RD}$ H382A** | (300-922), 62901 Da | - |
| **$AHK5_{RD}$** | (774-922), 16860 Da | Borkovcová et al., Phytochemistry 100 (2014) 6–15 |
| **$AHK5_{RD}$ D828A** | (774-922), 16455 Da | - |
| **$CKI1_{RD}$** | (944-1122), 15880 Da | Borkovcová et al., Phytochemistry 100 (2014) 6–15 |
| **$AHK4_{RD}$** | (944-1080), 15265 Da | Borkovcová et al., Phytochemistry 100 (2014) 6–15 |
| **AHP1** | (1-154), 22617 Da | - |
| **AHP2** | (1-156), 22341 Da | Degtjarik et al., Acta Cryst. (2013). F69, 158–161, Pekarova et al., Plant Journal (2011) 67:827-839 |
| **AHP3** | (1-155), 22344 Da | Pekarova et al., Plant Journal (2011) 67:827-839 |
